# Remote sensing of endogenous pigmentation by inducible synthetic circuits in grasses

**DOI:** 10.1101/2025.06.20.660755

**Authors:** Dong-Yeon Lee, Lucia Acosta-Gamboa, Luke Saleh, Sunita Pathak, Nathan Swyers, Austin Morgan, Susan Meerdink, Carolyn Kuzio, Santiago Calderon, Hudanyun Sheng, Samuel Kenney, Alina Zare, Malia Gehan, Dmitri A. Nusinow

## Abstract

Plant synthetic biology holds great promise for engineering plants to meet future demands. Genetic circuits are being designed, built, and tested in plants to demonstrate proof of concept. However, developing these components in monocots, which the world relies on for grain, lags behind dicot models, such as Arabidopsis thaliana and Nicotiana benthamiana. Here, we show the successful adaptation of a ligand-inducible sensor to activate an endogenous anthocyanin pathway in the C4 monocot model Setaria viridis. We identify two transcription factors sufficient to induce endogenous anthocyanin production in S. viridis protoplasts and whole plants in a constitutive or ligand-inducible manner. We also test multiple ligands to overcome physical barriers to ligand uptake, identifying triamcinolone acetonide (TA) as a highly potent inducer of this system. Using hyperspectral imaging and a discriminative target characterization method in a near-remote configuration, we can non-destructively detect anthocyanin production in leaves in response to ligands. This work demonstrates the use of inducible expression systems in monocots to manipulate endogenous pathways, stimulating plants to overproduce secondary metabolites with value to human health. Applying inducible pigmentation coupled with sensitive detection algorithms could enable crop plants to report on the status of field contamination or detect undesirable chemicals impacting agriculture, ushering in an era of agriculture-based sensor systems.

**Summary:** *Synbio tools for C4 grass model:*

● Advantage of synthetic switches as tools for biopharming and functional genomics
● Our workflow to optimize the gene circuits from a transient system to a stable transgenic
● Testing and taming golden gate elements in monocot system, *S. viridis*

## Introduction

Synthetic biology aims to improve genetic engineering tools to predictably control genetically encoded biological systems [1,2]. Despite recent advances in plant synthetic engineering, the development of tool kits and their supporting components is mainly confined to eudicots [3,4]. Adapting the existing synthetic biology tools directly to monocot grasses is challenging since monocots diverge from model eudicots in their promoter structure, cis-elements, and nucleotide composition [5,6]. Since deploying these tools will require significant optimization of components for use in monocots, the iterative Design-Build-Test (DBT) cycle of rational design in monocot model species will aid in facilitating gene discovery and improving agronomic traits in maize, rice, sorghum, and other highly productive food and bioenergy crops [7,8].

*Setaria viridis* (*S. viridis*) is a model for dissecting C4 photosynthesis and development in panicoid grasses [9]. Its short life cycle, small diploid genome, and transformability make it an ideal platform for studying the underlying mechanistic bases of agronomic traits in the closely related domesticated species *Setaria italica*, cereals (sorghum and corn), and bioenergy crops (sugarcane and switchgrass) [10]. Despite serving as a model for panicoid grasses, the available genetic toolkits for *S. viridis* have been limited to ectopic expression, RNA interference (RNAi), and genome editing [11,12]. Inducible promoter systems are desirable when constitutive expression of a transgene is likely to compromise plant development or metabolism [13]. Recently, the pOp6/LhGR system has been adapted to using two heterologous reporter genes, β-glucuronidase (GUS or *UidA*) and a yellow fluorescence protein (*YFP*) for use in rice [14,15]. Two chimeric components, a *pOp6* (6x lac operators and minimal *CaMV 35S* promoter) and an LHGR (fusion protein of *E.coli lacI* DNA binding domain, yeast Gal4 trans-activation domain ii (GAL4-ii), and rat glucocorticoid receptor (GR)) function as an inducible transcription system through the sequestration of the GR ligand-binding domain by chaperones until the presence of the steroid ligand dexamethasone causes folding and release [16]. We sought to determine if this system could be adapted for use in *Setaria* to drive the expression of exogenous reporters or transcription factors to alter endogenous pigmentation pathways to monitor changes in gene expression non-invasively [17,18].

Anthocyanin is a sub-branch of flavonoids and pigments that confers a broad spectrum of colors from orange, red, and purple to blue in reproductive and vegetative organs in plants [19], [20]. The biosynthesis of anthocyanins and flavonoids in plants is generally associated with abiotic stress responses, especially UV radiation, salt, and drought stress (reviewed in [21]). For humans, diets containing anthocyanin-rich fruits and vegetables are promoted for their antioxidant properties, which are linked to the prevention of age-related chronic disease and cancer and are a human health benefit [22,23]. The conspicuous coloration of anthocyanins has also served as a trait to dissect the genetic bases of the biosynthetic pathway underlying maize kernel pigmentation [24]. Combinatorial presence of both R2R3-MYB domain anthocyanin regulator *colorless aleurone* (*C1* or its paralog *PL1*) and basic helix-loop-helix proteins, *Colored 1* [*R* or its paralog *Booster* (*B*)] control the expression of anthocyanin biosynthetic pathway genes (CHALCONE SYNTHASE, CHALCONE ISOMERASE, FLAVONOID 3-HYDROXYLASE, DIHYDROFLAVONOL 4-REDUCTASE, ANTHOCYANIN SYNTHASE and UDP-GLUCOSYL TRANSFERASE) in a spatiotemporal manner in maize [25,26]. C1/Pl and R/B cloning in maize identified orthologous genes in many other species via a heterologous hybridization and sequence homology [26]. Ectopic expression of both R and C1 homologs simultaneously produces anthocyanin in several plant species, including tomato, maize, and rice [27–29]. Here, we sought to identify the native R and C1 orthologs in *Setaria viridis* and test their effectiveness in producing anthocyanins in both mesophyll protoplasts and stable transformants. We demonstrate that anthocyanin pigmentation can be a robust marker for testing programmable genetic circuits in monocot cereal species.

We show that constitutive expression of Sv*R1* and Sv*C1* results in high anthocyanin production throughout *S. viridis* and visibly purple plants. However, inducible anthocyanin pigmentation results in localized anthocyanin production, which is more challenging to detect. Traditional, destructive methods of anthocyanin quantification can detect low quantities of anthocyanin but require mechanical disruption of plant tissue, chemical extraction, and spectrophotometry to calculate anthocyanin abundance [30].

Hyperspectral imaging is a non-destructive method of measuring spectral information in a spatial context, and established vegetative indices have been used to estimate anthocyanin content [31–33]. Here, we utilize hyperspectral imaging and compare established vegetative indices for anthocyanin detection with potentially more sensitive discriminative target characterization and detection methods [34] for near-remote detection of anthocyanin production in *S. viridis* leaves. Coupling ligand-dependent pigmentation with remote detection could enable the development of sentinel plants that detect and report on the presence of chemicals in various agricultural settings.

## Results

### Identification of SvR1 and SvC1 in *S.viridis*

To identify the regulatory genes in the anthocyanin pathway in *S. viridis*, we constructed a phylogenetic tree of bHLH transcription factors related to known modulators of anthocyanin production in both monocots and dicots, including *ZmR*, *PhAN1*, *AtTT8*, *PhJAF13*, and *AmDEL* from *S. italica*, *S. viridis,* and other homologs in monocot species [35–37] (Fig. S1b). Sevir.5G416900 and Sevir.7G207500 from *S. viridis* are located in the same clades with *ZmR*, *ZmB,* and two R1-related genes in *S. italica* (Fig. S1a). We cloned Sevir.7G207500.1 using RT-PCR from extracted RNA of an ABA-treated leaf from the *Me034v-1* accession using primers spanning from the 5’UTR to the 3’UTR of the annotated gene (Fig. S1a, materials and methods). Multiple clones were identical to Sevir.7G207500.1, the putative SvR1 locus, from the *S. viridis* (A10.1) reference sequence, except for two single-nucleotide polymorphisms (SNPs) (Fig. S1c). Two additional cDNAs cloned corresponded to *Sevir.5G416900,* which shows 96% nucleotide identity to *SvR1*.

To locate the orthologs to *ZmC1*, we surveyed the C1 homologs in both *S. viridis* and *S. italica*. A phylogenetic tree was constructed with the homologs of the ZmC1 MYB transcription factor from *S. viridis, S. italica*, rice, and maize. Two clades formed into monocot-specific sub-clades, a C1 and P1 clade, and dicot homologs such as *AtPAP1*, *PhAN2*, and *AmRosea*, form separate branches at the other ends (Fig. S2D) [37–39].

The closest gene to Zm*C1* in *S. italica, Seita.4G086300,* contained a premature stop codon that would produce a truncated protein (Fig. S2B). C1 was unannotated in the *S. viridis A10.1* genome and thus missing in the phylogenetic tree of the C1-related MYB gene family (Fig. S2D). Manual inspection of syntenic regions in the *A10.1* genome to *SiC1* allowed us to identify a candidate full-length coding sequence homologous to *ZmC1* and *OsC1* (Fig. S2A and C). Interestingly, we found the presence of a *copia20* LTR transposon inserted at the C’ terminus *C1* region in the sequenced *Yugu1 italica* accession (Fig. S2A) [40]. Like other canonical copia-type transposons, the 3’ LTR end of the transposon introduced a stop codon that would lead to early translational termination of SiC1 (Fig. S2C), and 5 bp target site duplications were observed at the flanking site of transposon insertion [41]. The 5 kb long *copia 20-like* transposon is also present in the *B100 S. italica* accession, but not in *S. viridis* accession *Me034v-1* (Fig. S2B). We synthesized a golden gate module corresponding to SvC1 cDNA based on the sequence of *SvC1* regions in *Me034v-1* (Fig. S2B and Table S1).

### Ectopic expression of SvR1 and C1 together in mesophyll protoplasts is sufficient for anthocyanin production

To explore if the role of SvR1 and SvC1 in the promotion of anthocyanin biosynthesis was conserved in *S. viridis*, we made a series of constructs to express the transcription factors under the control of constitutive promoters in *S. viridis* protoplasts (Fig. 1A).

**Figure 1.**
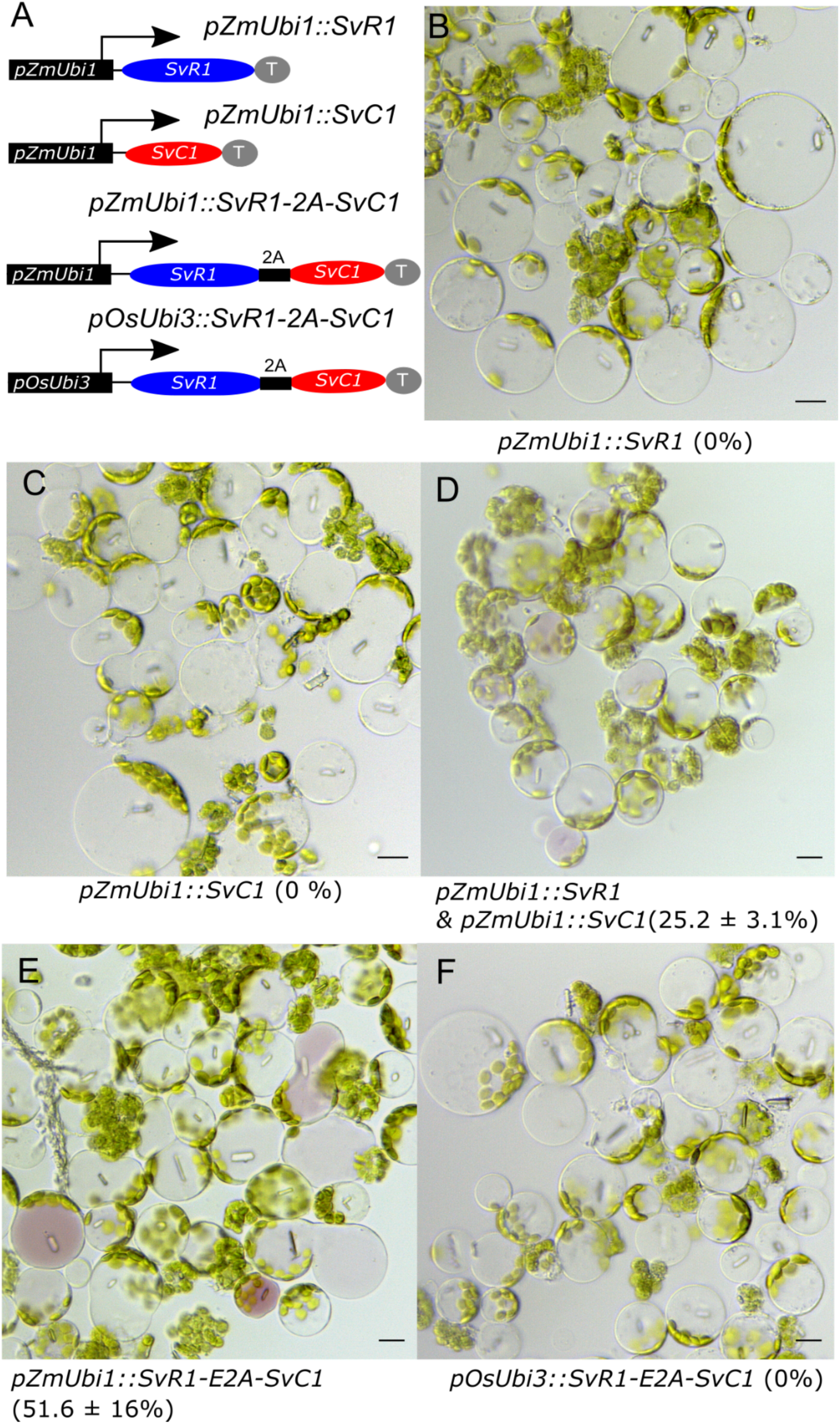
*SvR1* and *SvC1* are sufficient to induce anthocyanin production in *S. viridis* protoplasts. (A) Schematic representation of constructs transfected in *S. viridis* protoplasts. 2A, 2A skipping peptide; T, transcriptional terminator; *pZmUbi1*, *Zea mays Ubiquitin1* promoter; *pOsUbi3*, *Oryza sativa Ubiquitin3* promoter. (B-F) Microscopic images of the protoplast were taken at 50 hours after transfection with the constructs. *pZmUbi1::SvR1* (B), *pZmUbi1::SvC1* (C), *pZmUbi1::SvR1* & *pZmUbi1::SvC1* (D), *pZmUbi1::SvR1-2A-SvC1* (E) and *pOsUbi3::SvR1-2A-SvC1* (F). Numbers in parenthesis indicate the frequency of protoplast showing anthocyanin pigmentation from 2 independent transformations. At least 110 protoplasts per transfection were counted. Scale bar = 10 μm.

Ectopic expression of either *SvR1* or *SvC1* alone did not induce any noticeable changes in *S. viridis* mesophyll protoplast pigmentation (Fig. 1B and C). However, ectopic expression of both *R1* and *C1* transcription factors by co-transforming each construct together is sufficient to induce anthocyanin biosynthesis and pigmentation in protoplasts within two days (Fig. 1D). In addition, we made multigene *SvR1-E2A-SvC1* constructs, in which a 2A self-cleaving peptide sequence coupled to a glycine-serine-glycine spacer (GSG) was inserted between *SvR1* and *SvC1* for translating both proteins in equimolar amounts from a single promoter [42]. Compared to protoplasts transformed simultaneously with the two separate constructs, transformation with the *pZmUbi1::SvR1-E2A-SvC1* construct increased anthocyanin accumulation and nearly doubled the frequency of protoplasts with pigmentation (Fig. 1D & E). However, high expression of R1 and C1 is required for anthocyanin production, expression of *R1-E2A-C1* cassette from a weaker *Oryza sativa Ubiquitin 3* promoter (pOsUbi3) did not induce pigmentation of transformed protoplasts (Fig. 1F). Our protoplast transfection study demonstrated that both R1 and C1 are required for increasing pigmentation in *Setaria* protoplasts and that a single multigenic construct is sufficient to function as a reporter.

### Systemic anthocyanin production in *S. viridis* by constitutive expression of *SvC1* **and *SvR1***

To test if the expression of *SvC1* and *SvR1* can induce pigmentation in *S. viridis* plants, we assembled two *SvR1-E2A-SvC1* multigenic L1 constructs expressed from either the *ZmUbi1* or *OsUbi3* promoter together with an L1-hygromycin resistance cassette into L2 binary constructs for plant transformation (Fig. 2A). Introduction of pSENT98 (pZmUbi1::SvR1-E2A-SvC1) induced purple pigmentation from the callus during tissue culture (Fig. S3A) and many of the pSENT98-transformed *Setaria* seedlings displayed a broad spectrum of purple coloration ranging from mosaic to uniform (Fig. S3C-D). T0 regenerates with deep purple coloration tended to be compromised in their growth and fertility (Fig. S3C). However, we were able to isolate a stably introgressed purple-hued *Setaria* transgenic from one pSENT98 event, which showed the heterogeneity in transgene structure at primary generation and subsequently segregated into high-copy transgenic lines and a single copy at T1 generation based on our Taqman based genotyping analysis (Fig. 2B and S3D, data not shown). Fortunately, the single copy line from pSENT98 #1 showed stable anthocyanin pigmentation throughout its life cycle and for multiple generations. However, pSENT99 (pOsUbi3::SvR1-E2A-SvC1) transgenics only showed noticeable color changes in the ligule region and senescing leaves after heading (Fig. S3G-I). Other multi-T-DNA insertional pSENT98 events displayed varied anthocyanin pigmentation, which correlated to the extractable amount of anthocyanin as measured by spectrophotometer (Fig. S3D, E). A tandem 6xHis-and Flag-epitope tag (HFC) at the C-terminus of SvC1 in both pSENT98 and pSENT98 was used to compare the protein expression level and 2A peptide self-cleaving efficiency (Fig. 2C). SvC1-HFC protein level correlated with the anthocyanin pigmentation in pSENT98 leaf blades, but undetectable in leaf blades of pSENT99 #12 plants, corroborating the lower activity of OsUbi3 promoter (Fig. 2C). This low activity of the OsUbi3 promoter contrasted with the rice study, in which the same *OsUbi3* promoter of pRESQ48 showed a 2.2-fold higher induction compared to the *ZmUbi1* promoter in rice suspension cell study [43]. In addition, the high efficiency of self-cleaving by our 2A skipping peptide design in the golden gate system was confirmed by the lack of any uncleaved polypeptides of SvC1-HFC in longer exposure (data not shown). Altogether, our transgenic approaches demonstrate that anthocyanin biosynthesis in *S. viridis* by our *SvR1-E2A-SvC1* reporter is an effective non-invasive visual marker to evaluate promoter activity with the naked eye.

**Figure 2.**
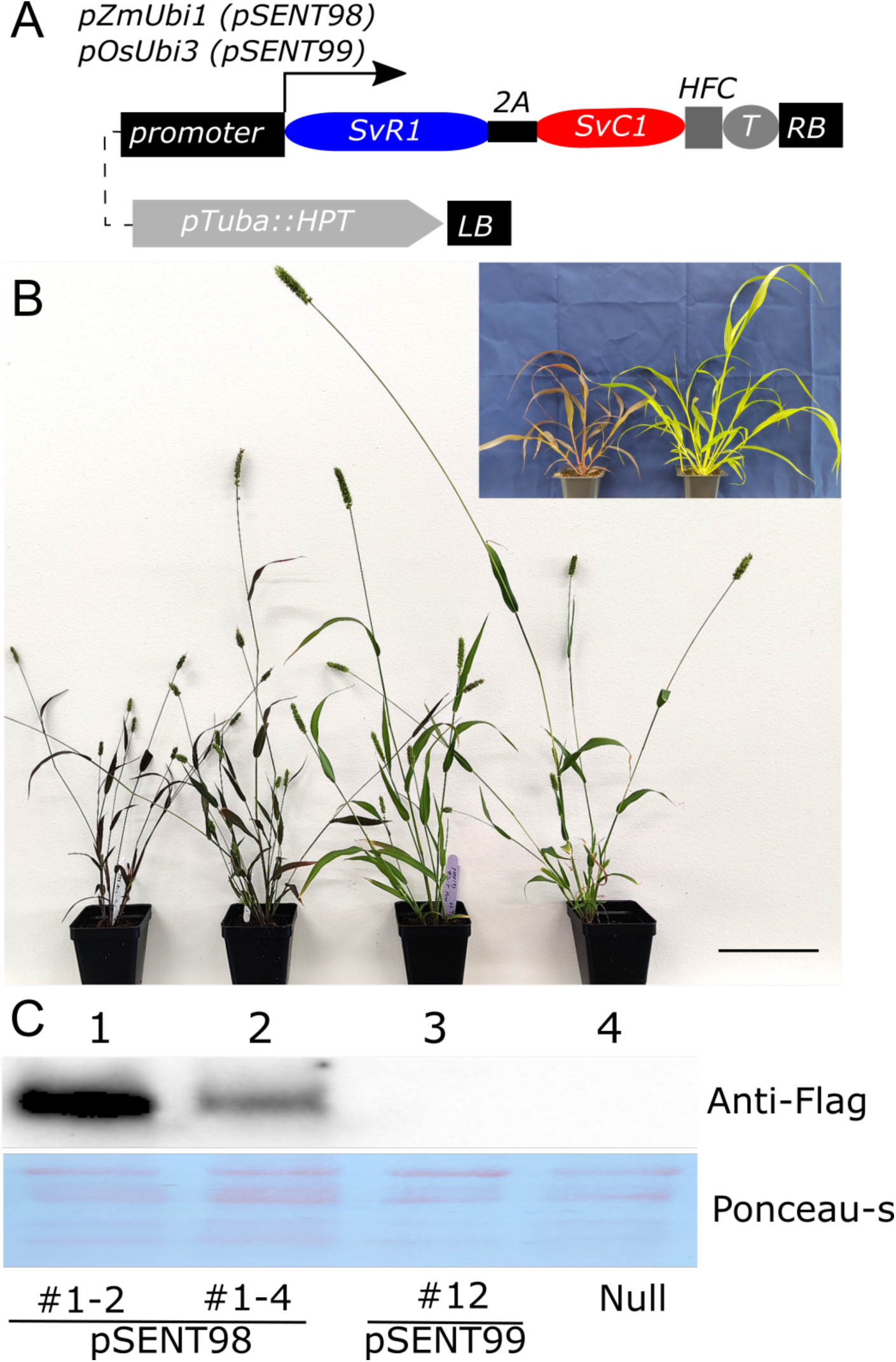
Simultaneous expression of *SvR1* and *SvC1* induce the anthocyanin production *in vivo*. (A) Construct layouts of pSENT98 and pSENT99 for stable transformation. pTuba::HPT, hygromycin resistance gene cassette (pOsTub1a::Hygromycin phosphotransferase:OsTub1aT); LB, Left border of T-DNA; RB, Right border of T-DNA; HFC, C-terminal 6x His-and 3x Flag tag; T, terminator. The arrow indicates the direction of transcription. The filled box and ellipses represent promoter and gene coding sequences, respectively. (B) Anthocyanin phenotype of representative T3 plants of pSENT98 #1-2, pSENT98 #1-4, pSENT99 #12, and null segregant, from left to right, in the same order of western blot lanes at 21 days after sowing, scale bar=10 cm. Inset image shows the hue of anthocyanin plant (pSENT98 #1-2,left) and its null (right). (C) Protein expression of SvC1-HFC in pSENT98 and pSENT99 transgenic lines and controls from leaves of 10-day old plants. Anti-FLAG antibody was used to detect SvC1-HFC (C-terminally fused His-FLAG tag). Lane 1, pSENT98 #1-2; 2,pSENT98 #1-4; 3, pSENT99 #12; 4, Null, in the same order of plants in (B).

### Rewiring a synthetic circuit to regulate the endogenous anthocyanin pathway

Having shown that the *SvR1-E2A-SvC1* cassette was a promising visual marker when driven from constitutive promoters, we wanted to determine if it could be used to monitor changes in promoter activity. A 6x Lac Operator minimal 35S promoter fusion (pOp6) from the *rice codon optimized LHGR-N* (*GR-LacI-Gal4-ii,* domain fusion in order)/*pOp6* dexamethasone-inducible circuit was assembled with *SvR1-2A-SvC1* (Fig. 3A) [15]. In addition to *R1* and *C1*, we also incorporated a *Pyrearinus termitilluminans* larval click beetle luciferase (*ELUC*) into the SvR1-2A-SvC1 cassette linked by an additional P2A skipping peptide for real-time monitoring of bioluminescence [44] (Fig. 3A and Table S1). As an effector construct, a rice-codon-optimized *LHGR-N* fusion gene paired with a strong *ZmUbi1* promoter or weak *Zea mays Elongation factor 1a* (*ZmEf1a*) promoter, which shows a 30 ∼ 70 fold difference in the expression of the luciferase reporter in *S.viridis* protoplasts (Fig. S4) [15]. Additionally, the *Drosophila Gypsy insulator* was placed before the *pOp6* promoter to mitigate the possibility of an unknown enhancer activity from the plasmid backbone [45]. Luciferase expression of the triple gene reporters showed a rapid increase after adding dexamethasone (Dex) that peaked at 6-7 hours and slowly declined thereafter (Fig. 3B). In comparison, *pZmEF1a::LHGR-N* reduced the basal expression of luciferase and resulted in a 6.3 fold induction (six hours after addition of 10 μM Dex), compared to the 3.4 fold induction from a *pZmUbi1* driven effector at a similar time point. However, the strong LHGR-N expression by *pZmUbi1* produced nearly twice the maximum luminescence compared to transfections with *pZmEf1a::LHGR-N*, but showed higher background in the absence of Dex (Fig. 3B). Similar to our observations with luciferase expression, anthocyanin production was apparent in mock-treated samples regardless of promoter strength and increased two days after addition of Dex (Fig. 3C-D). The high levels of basal activity could be due to the excessive amounts of LHGR-N, which could surpass the level of endogenous proteins that sequester the GR receptor, by the strong *pZmUbi1* promoter, or by the high copy number of plasmids during PEG-mediated transfection.

**Figure 3.**
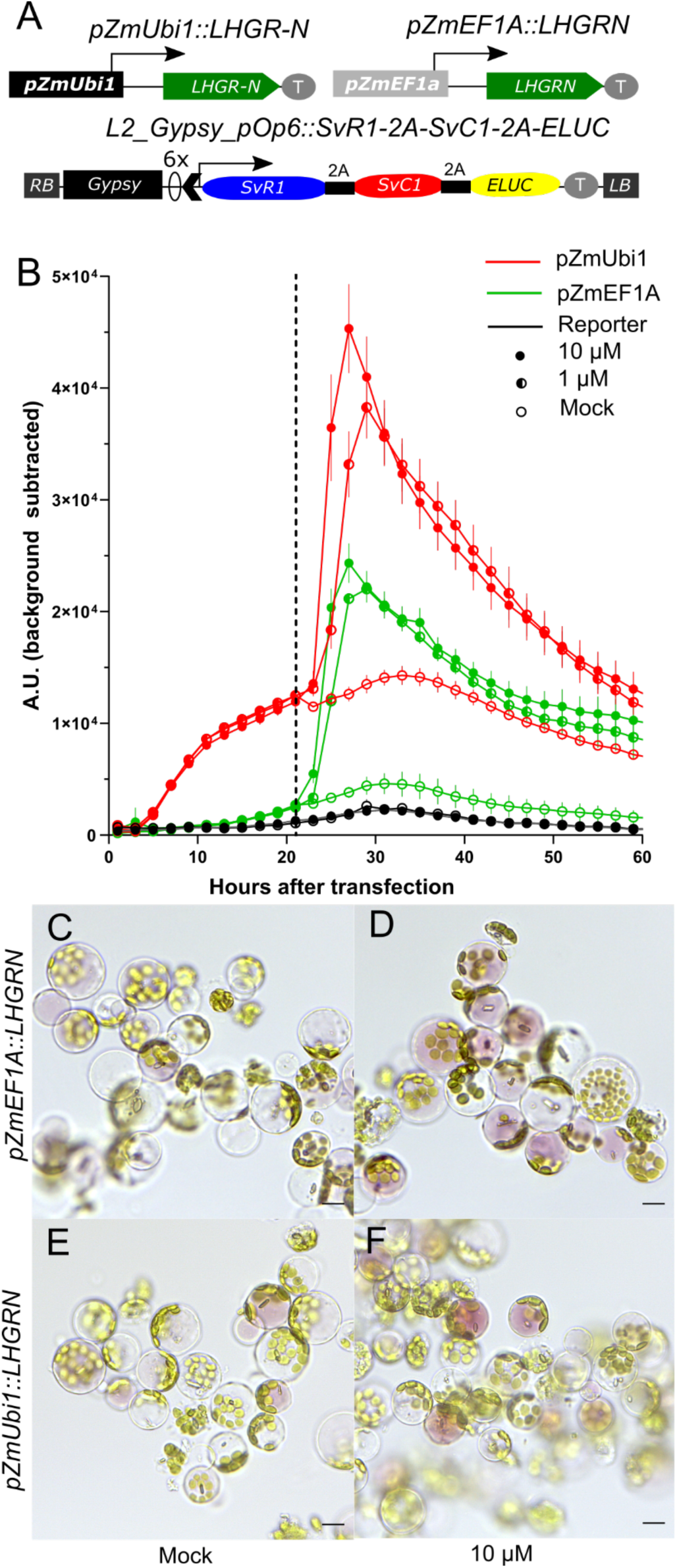
LHGR-N/pOp6 circuit induces the bioluminescence and anthocyanin with the presence of Dexamethasone in *S.viridis* protoplasts. (A) Construct schemes of dexamethasone inducible systems for protoplast transformation. *Zea mays* Ubiquitin 1 promoter and EF1a promoter were fused to the LHGR-N effector gene (top) and 6 x Lac Operator promoter (pOp6) was fused to the triple gene reporter (SvR1-2A-SvC1-2A-ELUC) in L2 construct. Gypsy, Drosophila Gypsy insulator; 2A, 2A-skipping peptide sequence; ELUC, *Pyrearinus termitilluminans* larval click beetle luciferase; RB, Right border of transfer DNA (T-DNA); LB, Left border of t-DNA; black chevron, minimal 35S promoter; O, Lac operator sequence. (B) Time course quantification of bioluminescence induction by dexamethasone in protoplast transfected with reporter and effector construct. pZmUbi1::LHGR-N (Red line) and pZmEF1A::LHGR-N (Blue line) and without effector construct (black line). Filled, half and empty circles indicate 10 μM, 1 μM dexamethasone (Dex) and mock (0.1% Met-OH) added to the respective well. The dashed line is the time point when Dex was added to each well. Each line is an average of two bio reps and the vertical line represents the standard error. A.U.; arbitrary unit of relative luminescence. (C-F) Dexamethasone inducible anthocyanin in the protoplast with LHGR-N/pOp6 circuits. *pZmEF1::LHGR-N* (C-D) and *pZmUbi1::LHGR-N* (E-F). Images were taken at 60 hours after mock treated (C, E) and 10 µM Dex treated (D, F), respectively. Scale bar = 10 µm.

To determine how these two promoter-driven circuits respond *in vivo* as stable transgenic, L2 constructs carrying both L1-LHGRN constructs were introduced to *S. viridis* Me034*v-1* (Fig. 4A and Fig. S5A). Primary regenerates of pSENT166 and pSENT162 didn’t show any noticeable phenotypes, unlike the pSENT98 transformants. Three independent single T-DNA insertional events secured from pSENT166 (pZmUbi1) displayed a dramatic induction of bioluminescence and peaked at around 35 hours after Dex application in our leaf-punch bioluminescence (BL) assay (Fig. 4B). Luciferase induction level was responsive to DEX concentration, saturating at around 1 μM concentration and showed basal bioluminescence in mock-treated samples, similar to the protoplast results (Fig. 4B). However, pSENT162 (pZmEf1a) transgenic events produced only around 40 fold lower bioluminescence induction compared to the pSENT166, but its luciferase induction correlated with Dex concentration (Fig. S5). Similar to bioluminescence induction (Fig. 5A), anthocyanin pigmentation was mostly limited to the marginal area or cut region of pSENT166 leaves by dipping in 10 μM Dex solution, indicating low perfusion of Dex through the leaf epidermal layer (Fig. 4C-D). To improve Dex diffusion into the tissue, various application methods (e.g., aerosol spray, leaf dipping, or watering) were tried with and without surfactants (Silwet L-77 or Break-Thru OE 446). Still, anthocyanin induction in leaves was inconsistent, both visually and by extraction (data not shown). We hypothesized that Dex may not be able to perfuse through the epidermis except in areas disrupted by cutting. To test this, we punctured tissue using a microneedle roller as described in a citrus transformation protocol [46] and exposed microperforated leaf tissue to Dex. The pigmentation pattern around the micro-perforated holes was clearly observed in the detached leaf dipped in Dex solution (Fig. 4C-D), but not in vehicle-treated control plants (Fig. 4E), showing that the pigmentation was not a response to the physical damage. This experiment showed that low penetration of dexamethasone through *S. viridis* leaf tissue likely impeded induction of the circuit. To improve the Dex infusion *in planta* of *S. viridis*, we tried ultrasonic fogging of the Dex solution by nebulization [47]. Overnight nebulization of pSENT166 transgenic lines produced more consistent anthocyanin induction in colorimetric extraction, but the leaf color change was still limited in the marginal area of the exposed tissues (Fig. 5E).

**Figure 4.**
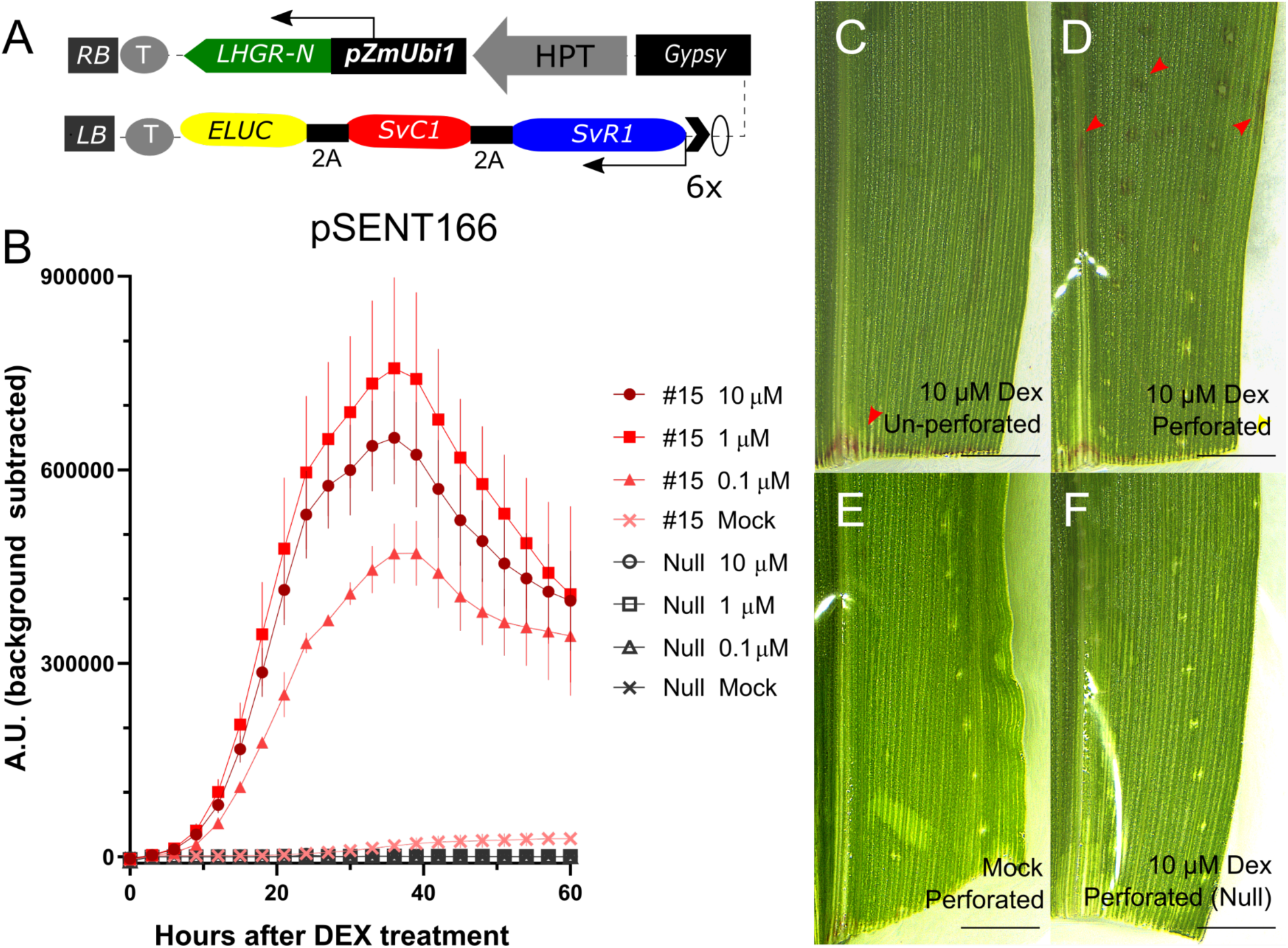
LHGR-N/pOp6 circuit induces the bioluminescence and anthocyanin with the presence of Dexamethasone in stable *S.viridis* transgenics. (A) Construct layout of Dex-inducible stable constructs, pSENT166. HPT, hygromycin resistance transcription unit. Gypsy, Drosophila Gypsy insulator; 2A, 2A-skipping peptide sequence; ELUC, *Pyrearinus termitilluminans* larval click beetle luciferase; RB, Right border of transfer DNA (T-DNA); LB, Left border of t-DNA; black chevron, minimal 35S promoter; O, Lac operator sequence. (B) Bioluminescence induction from pSENT166 #15 leaf discs in response to a serial dilution of dexamethasone. Red and grey lines represent homozygous and nulls, respectively. Vertical line of each point indicates the standard error for 3 bio-reps from individual plants. (C-F) Anthocyanin induction to Dex limited to the marginal and wound region of pSENT166 leaf. Leaf blades were submerged to the 10 μM dexamethasone-containing water (C, D and F) or Mock (0.1% Methanol, E) with 0.001% BT OE044 for 2 days. Leaf blades cut from T2 transgenic, pSENT166 #8 T2 (D - E) and their null (F) were used to be microperforated using a derma-roller before incubation to enhance the ligand penetration into leaf tissue. Red arrowhead pointed out the anthocyanin pigmentation around leaf cuts, punctures and marginal areas etc. Scale bar=1 mm.

**Figure 5.**
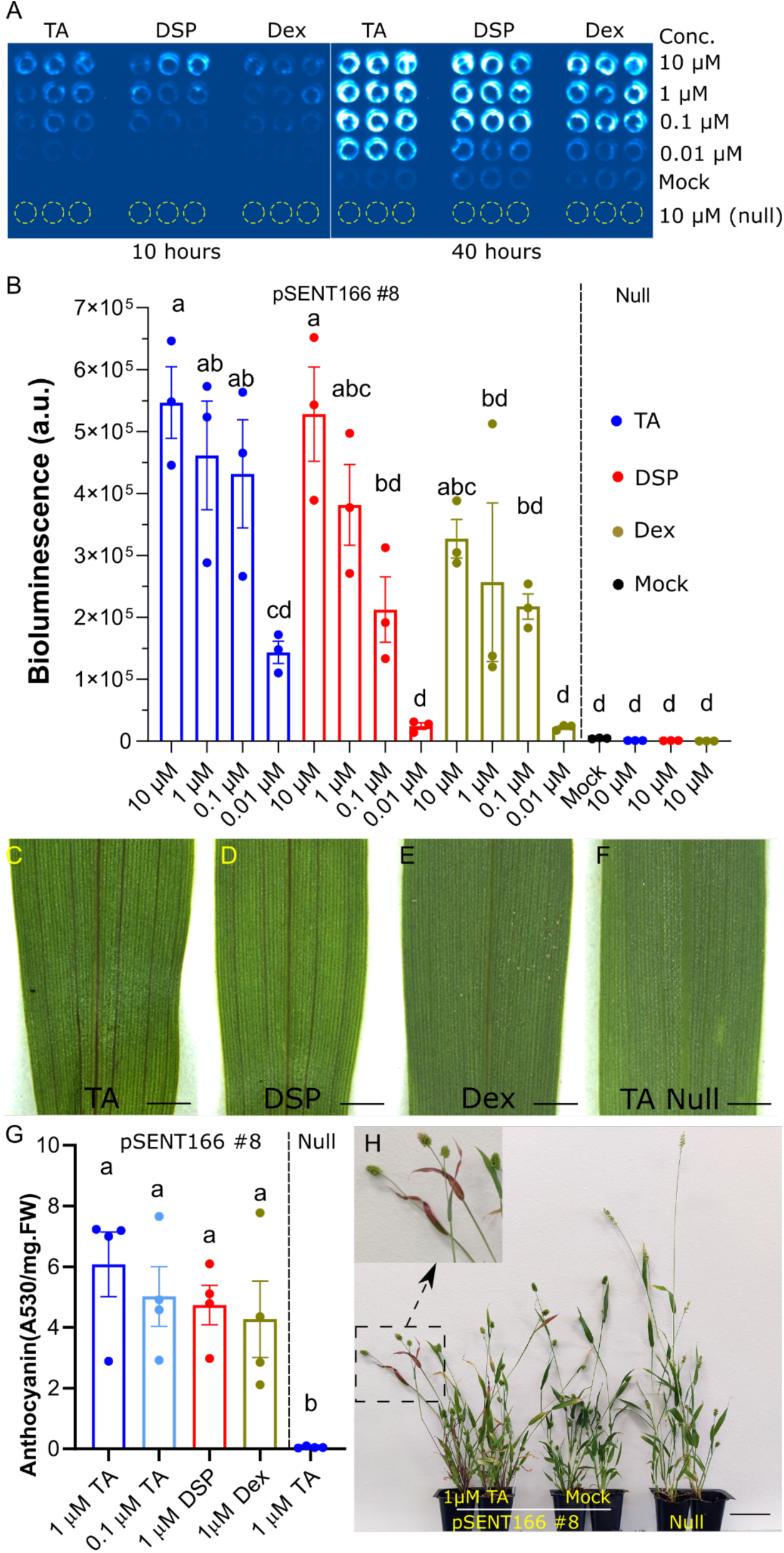
Triamcinolone Acetonide and Dexamethasone sodium phosphate increase LHGR-N/pOp6 driven reporter expression. (A) Bioluminescence induction of pSEN166 to the Dex analogs, Triamcinolone Acetonide (TA), Dexamethasone sodium phosphate (DSP) and Dexamethasone (Dex) were added to each well as indicated concentration at the right of image. Image of 2 time points are shown as representatives of steroids diffusion into leaf discs. Leaf discs from 3 individual transgenic and null are laid out and nulls were highlighted with yellow dashed outlines due to their absence of bioluminescence. (B) Dose dependent bioluminescence induction (arbitrary unit (A.U.)) with the various concentrations of different steroids at 20 hours after treatment. Data presented are mean ± standard errors of three replicates. TA, DSP, Dex and Mock are separated by blue, red, brown and black color of bar and their concentration is represented on the x-axis. Bars with different letters are significantly different by Tukey’s post-hoc one way ANOVA analysis (one-sided) (P<0.05). (C-F) anthocyanin accumulation along vasculature shown on the leaf blade of pSENT166 treated by the different Dex analogs. Whole plants are fumigated overnight with the indicated steroid species (bottom) in the sealed container. Images were taken 5 days after the nebulization of 1 μM steroid. Scale bar = 1mm. (G) Anthocyanin contents of pSENT166 nebulized with the different steroids. Data represents mean ± standard errors of 4 biological replicates (dots) at 5 days post treatment. Bars with the different letters are significantly different in Tukey’s post-hoc ANOVA analysis (P<0.05). (H) long-term anthocyanin induction of pSENT166 treated with the 1 uM TA diffusing. Close-up of red-pigmented flag leafs is shown in the inset. Images were taken 35 days after the treatment. Scale bar = 5cm

We considered that changing the ligand to one with better absorption properties might improve uptake and induction. [14] reported that triamcinolone acetonide (TA) can be a more potent steroid in rice than dexamethasone. We also included highly water-soluble dexamethasone sodium phosphate (DSP) in addition to TA and Dex to determine which steroid is more effective in turning on the reporter through *Setaria* leaf tissue. In our leaf-punch BL assays on pSENT166, 10 μM TA or 10 μM DSP induced bioluminescence faster (peaking at 20 hours) compared with 10 μM Dex at the same time point (Fig. 5A-B). A lower concentration of TA was more potent in bioluminescence induction compared to the same concentration of DSP and DEX (Fig. 5A-B), showing 5-6 fold induction at 0.01 μM TA compared to 0.01 µM DSP or Dex. Similar to the rice studies, 10 μM Dex resulted in lower maximum BL than 1μM Dex (Fig. 5B) and visual leaf damage, indicating possible toxicity (data not shown) [14,15]. Meanwhile, TA and DSP didn’t cause any compromise in BL at 10 μM and showed faster induction to maximum BL and more expanded BL to the center (Fig. 5A-B). Ultrasonic nebulization of pSENT166 plants with these steroids showed that 1µM TA nebulization led to higher anthocyanin accumulation than plants with the same dose of DSP and DEX (Fig. 5G).

Despite small differences in total anthocyanin content, TA and DSP nebulization developed a distinct anthocyanin pattern along the vasculature (Fig. 5C-D), indicating that TA and DSP penetrated more readily than Dex through *Setaria* leaf tissues at 1µM.

More importantly, TA nebulized plants showed anthocyanin accumulation in exposed tissue and in newly developed tissues, flag leaf, and internodes of panicle (Fig. 5H and S6). Our luciferase assay and anthocyanin accumulation showed that TA is the most effective steroid in reporter expression of *Setaria* LHGRN/pOp6 among the tested ligands, and ultrasonic nebulization is an effective chemical delivery method ‘*in planta*’.

### Hyperspectral Imaging-based Detection of Anthocyanin Generation

We anticipate that the imaging of sensor plants by Unmanned Aerial Vehicles (UAV) or aerial platforms would complement their deployment for field applications. Hyperspectral image data (details about sensor in Methods) were collected on wild-type *S. viridis*, accession *Me034v-1*, and pSENT98 transgenic constitutively expressing anthocyanin TFs, *SvC1* and *SvR1* (Fig. 2A-B) to assess if anthocyanin pigmentation can be detected through imaging. Two anthocyanin detection approaches were applied and compared.

The first approach relied on the Anthocyanin Reflective Index (ARI), a well established vegetation spectral index, to non-destructively quantify relative anthocyanin content [33]. PlantCV [48,49], an open-source and open-development image analysis software, was used to segment plants from the background in hyperspectral images (analysis workflows available here: https://github.com/danforthcenter/acosta-gamboa-anthocyanin). Two spectral bands, 550 ± 15 and 700 ± 7.5 nm, were used to calculate ARI [33] (Fig. 6). A histogram of the average ARI index is significantly different (KS Test =< 2.2e-16) between pSENT98 and wild-type (Fig. 6A). pSENT98 are also visibly purple to the naked eye compared to wild-type (Fig. 6B). This result demonstrates the feasibility of detecting anthocyanin using hyperspectral imaging at 0.3 meters and the ARI index in *S. viridis* plants constitutively expressing *SvC1* and *SvR1*.

**Figure 6.**
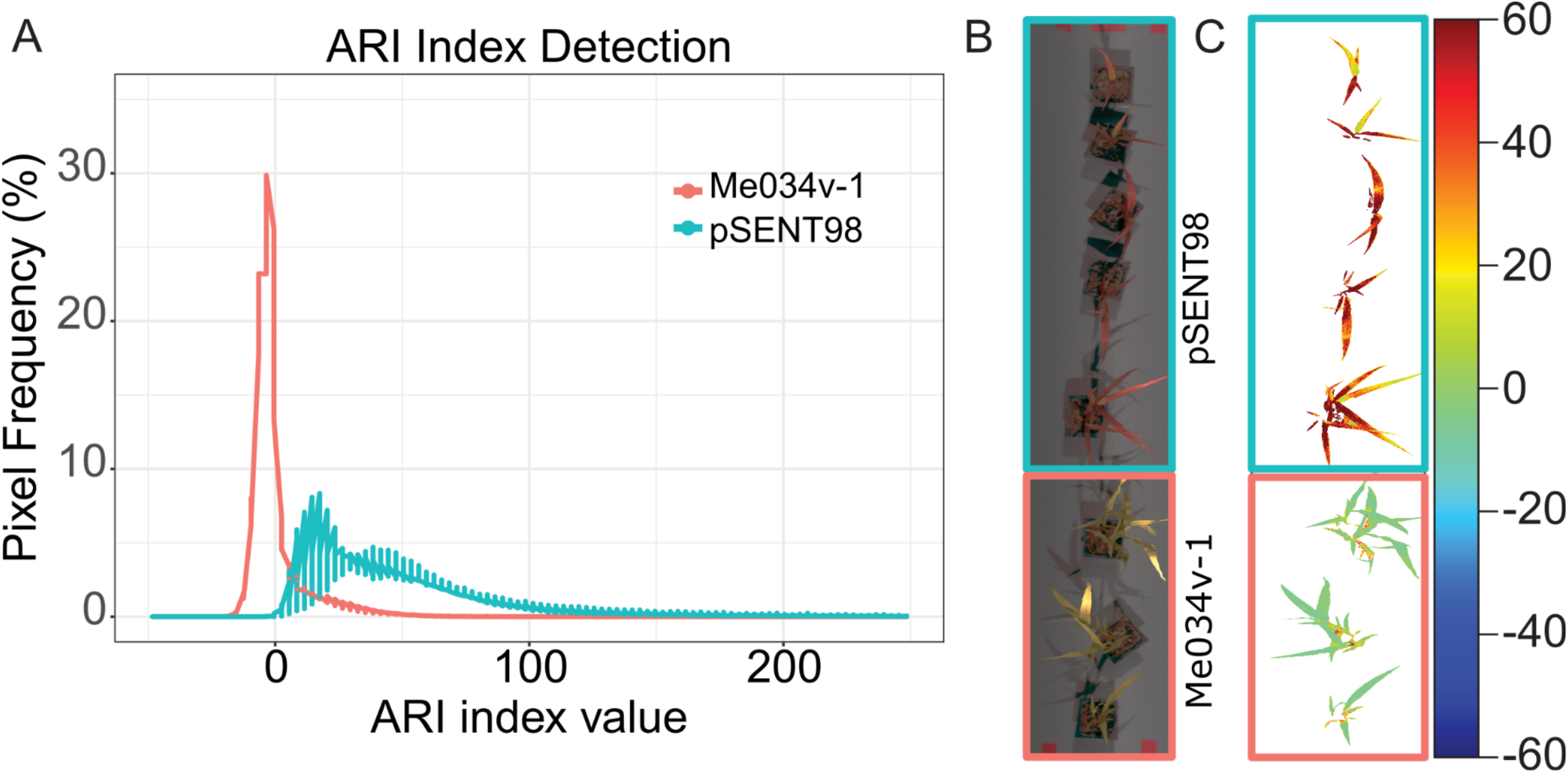
Detection of anthocyanin by ARI index in S. viridis constitutively expressing SvC1 and SvR1 (pSENT98). (A) Anthocyanin reflectance index (ARI) from plants constitutively expressing SvC1 and SvR1 (pSENT98) and WT setaria accession ME034v-1. The lines in the graph represent the average ARI index value, n=5 for overexpressing lines and n=3 for Me034v-1. KS test: p-value < 2.2e-16. (B) Pseudo RGB with pSENT98 plants boxed in blue and WT (ME034v-1) in red. (C) Pseudocolor images of ARI index values. pSENT98 are five plants boxed in blue and wild-type plants three plants boxed in red.

To demonstrate the feasibility of detecting ligand-induced anthocyanin production, which we expect would be much more spatially localized, we tested the ARI index on the Dex inducible LHGR-N/pOp6::R1-C1-ELuc line (pSENT166). Two independent pSENT166 events, #6 and #8, and a null sibling, were sprayed with TA, DEX, and DSP to determine if hyperspectral imaging can detect anthocyanin induction of the circuit throughout plant tissue. A statistically significant shift in ARI index values was detected between the transgene null sibling (pSENT166 #8) sprayed with TA and the pSENT166 #6 sprayed with TA (KS Test = 2.861e-08), DEX (KS Test = 0.007992), or DSP (KS Test = 0.04463). A statistically significant shift was also detected between the transgene null sibling (pSENT166 #8) sprayed with TA and pSENT166 #8 lines sprayed with TA (KS Test = 0.0001824), DEX (KS Test = 0.01008), or DSP (KS Test = 2.674e-05; Fig. S8B). Anthocyanin content quantified from leaves of the constitutive anthocyanin line (pSENT98) was approximately 73 A530/mg.FW (Fig. S3E), while the Dex inducible line pSENT166 treated with 1µM TA was around 6 anthocyanin (A530/mg.FW) (Fig. 5G).

Therefore, an ARI index can be used to discriminate *S. viridis* plants constitutively expressing anthocyanin (pSENT98) and *S. viridis* plants with induced expression of anthocyanin (pSENT166) from wild-type *S. viridis*. However, while the KS tests are statistically significant, this subtle shift is difficult to discern without statistical calculations, and the probability of a false alarm as calculated by a Receiver Operating Characteristic (ROC) curve is high and close to a random classifier (Fig. 7D). Therefore, we tested other image detection methods that might be more robust when expression of anthocyanin is lower and likely also more localized.

**Figure 7.**
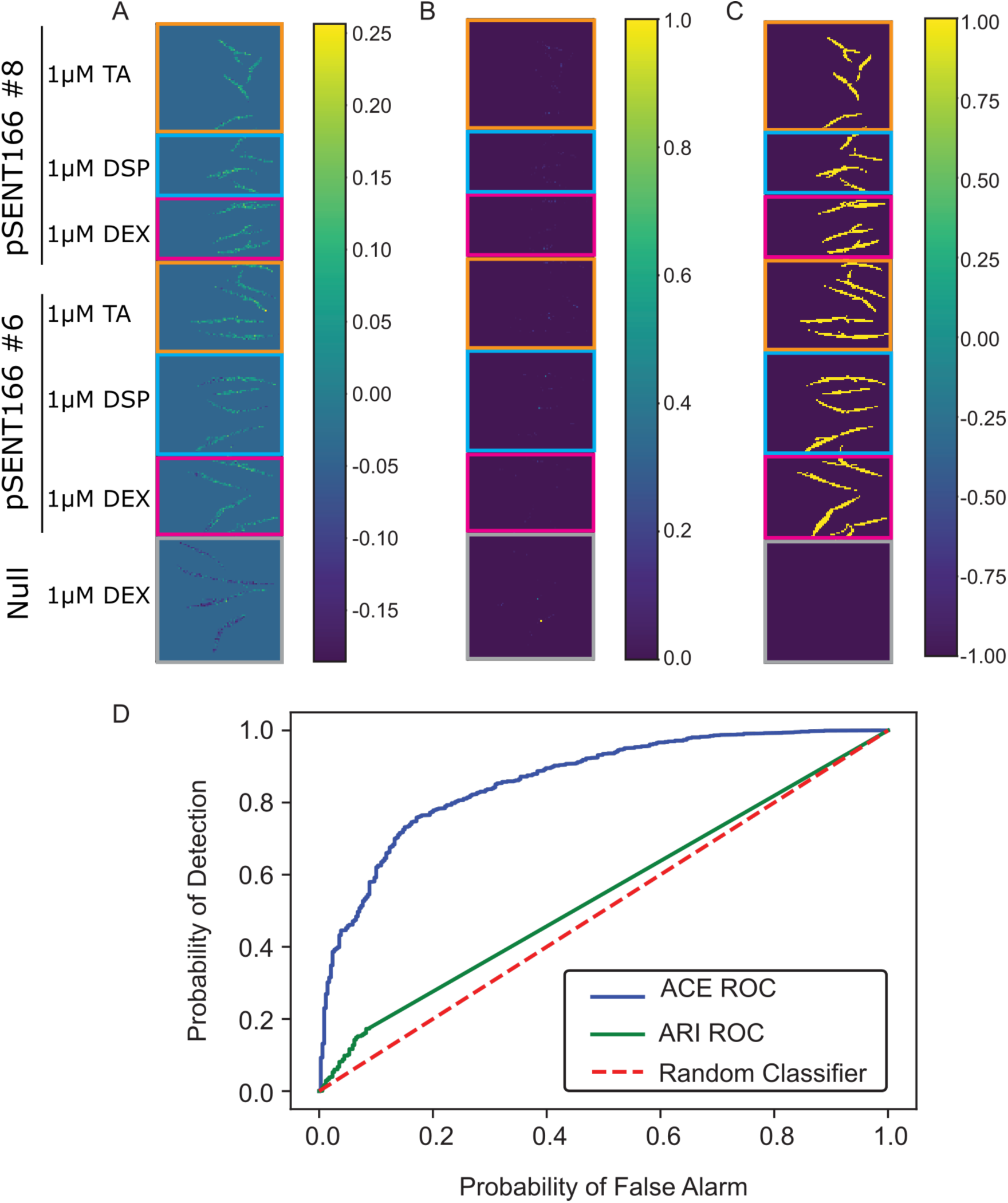
Application of Multiple Instance Adaptive Cosine Estimator (MI-ACE) to detect anthocyanin content in pSENT166 (#6-7 and #8-7) setaria plants as well as the null line (#8-10) sprayed with 1µM DSP, TA, or DEX. A) MIACE confidence map, B) ARI confidence map, C) Truth map (shows where anthocyanin was present, bright yellow means where anthocyanin is present, dark purple represents where anthocyanin is not present and blue corresponds to background. D) Receiver operating characteristic (ROC) curve for both ARI and MIACE approaches, MIACE shows a lower probability of false alarm and higher probability of detection.

The second detection approach relied on the Multiple Instance Adaptive Cosine Estimator (MI-ACE) approach for discriminative hyperspectral target characterization and detection. MI-ACE is a trained machine learning algorithm for sub-pixel target detection that learns a discriminative target representation that optimizes the Adaptive Cosine Estimator (ACE) [50] detection results on a training set even under conditions of imprecise labels. The MI-ACE algorithm is applied in two steps. The first step uses a training set labeled with target vs. non-target regions (in this application, regions with expected anthocyanin response vs. regions in hyperspectral imagery without any anthocyanin response). Using this training set, a discriminative target signature is iteratively learned and optimized using the methods described in the Materials and Methods section below. In the second step, the discriminative target signature can be used within the ACE detection measure along with the background mean and covariance estimated from the non-target training data to detect targets on unlabeled test imagery.

To develop the discriminative target signatures, each image is pre-processed to be split into multiple “bags,” or subsections (for analysis workflows see: https://github.com/GatorSense/MI-ACE-Notebooks). The data was split into its corresponding bags by visual inspection and division of image sections based on known locations of the anthocyanin presence. Bags were labeled as positive (contains at least one pixel with anthocyanin present) or negative (no anthocyanin present in the pixel). MI-ACE then optimized the Adaptive Cosine Estimator (ACE) detection statistic on this training data. As shown in Supplemental Figure 9, MI-ACE is very effective at matching expected anthocyanin locations (Supplemental Figure 9C) on a test data set from plants constitutively expressing anthocyanin (pSENT98; Supplemental Figure 9A). Furthermore, ROC analysis of the ARI and MI-ACE show comparable accuracy and sensitivity, with MI-ACE having a slight advantage in the probability of detection (Supplemental Figure 9D).

When we apply the MI-ACE algorithm to the detection of the inducible anthocyanin circuit, we find that MI-ACE vastly outperformed ARI in the ability to detect anthocyanin. ACE was able to distinguish the inducible anthocyanin lines pSENT166 #6 and #8 setaria plants sprayed with 1uM DSP, TA, and DEX from the null line (#8-10) sprayed with 1 µM TA (Figure 6A). ROC analysis on this dataset supports the superiority of MI-ACE over ARI for anthocyanin detection (Figure 7D).

## Discussion

These results show that circuits that alter endogenous pigmentation pathways in *S. viridis* can be used in conjunction with hyperspectral imaging to detect the presence of chemical ligands in the environment. Previously, an inducible LHGR/pOp6 system was successfully implemented in monocot rice using exogenous GUS and YFP reporters [14,15]. Here, we cloned anthocyanin pigmentation regulatory transcription factors, *SvR1* and *SvC1*, and show that ectopic expression of both transcription factors is sufficient to turn on the anthocyanin pathway in *S. viridis* mesophyll protoplasts and whole plants. Then, we developed a multi-gene reporter system coupled to LHGR-N/pOp6 modules and evaluated the reporter response against the steroids by multi-reporter imaging of mesophyll protoplasts and leaf discs from stable transgenic lines. Similar trends in anthocyanin expression using the designed constructs in both protoplasts and stable transgenic corroborated transient *S.viridis* protoplast transformation as an efficient platform to run the DBTL cycles for synthetic biology designs before moving to the more laborious process of stable transformation in a homologous system. Levering changes in leaf pigmentation due to altered anthocyanin absorption and reflectance, we used hyperspectral imaging and the Anthocyanin Reflective Index to evaluate and estimate anthocyanin content [51]. To overcome the non-uniform induction of the reporter in leaf tissue, we applied MI-ACE to develop a learning discriminative target concept for anthocyanin pigmentation. MI-ACE outperformed ARI on images from plants containing inducible circuits, while both methods performed similarly on plants constitutively expressing the reporter. Together, these results demonstrate a path for using synthetic biology to interface with endogenous pathways to alter plant pigmentation and hyperspectral imaging analysis to use plants as chemical sensors in the field. Below, we discuss some insights we have gained in developing these plant tools.

### Domestication of anthocyanin regulatory genes and transposon in selection in Setaria italica

To adopt the native anthocyanin biosynthesis as a marker in *S.viridis*, we isolated anthocyanin regulatory genes, *SvR1* and *SvC1*, using phylogenetic analysis and manual annotation. Interestingly, the lack of annotation corresponding to C1 in the *S. viridis* A10.1 genome and early translational termination of *SiC1* in the *S. italica Yugu1* genome led us to identify the presence of Copia-type transposon at the ’C’ terminus of *SiC1* gene. Copia-type Tn insertion in this loci is conserved in at least two *S. italica* accessions, *Yugu1*(*Yg1*) and *B100* (Fig. S2C). Recently, *SiR1* was pointed out as a candidate for the purple color of *pulvinus and leaf sheath* (*PPLS*), a key trait for screening hybrids in foxtail millet breeding, by association mapping between green sheathed *Yg1* and purple-sheathed *Shi-Li-Xiang* (*SLX*) [52]. Our finding of the presence of *copia20* LTR transposon in the C Terminus of *SiC1* in two landraces suggests that the truncated version of C1 may be associated with reduced anthocyanin pigmentation in certain *S. italica* landraces during domestication.

### Building the multigenic reporter for the inducible system

Previously, the LHGR/pOp6 system was successfully implemented in monocot rice using GUS reporter and YFP [14,15]. In addition to demonstrating the LHGR-N/pOp6 modules function in *Setaria*, we evaluated the reporter response against the steroids by the time-course luciferase imaging reporter in mesophyll protoplast and leaf discs from stable transgenic lines. The development of the reporter was facilitated by developing multigenic constructs compatible with the golden gate system by adding two 2A skipping peptides with the Glycine-Serine-Glycine linker at the N-terminus position of the L0-C1 position (Fig. S7a) [42]. The 2A peptide system allows for the expression of multicistronic genes from the same promoter and reduces promoter and terminator use, which are still limited in cereal species for plant biotechnology [53]. We generated multicistronic reporters to induce anthocyanin expression and incorporated a bright larval click beetle luciferase (ELUC) for monitoring bioluminescence [54]. Adding Eluc to the anthocyanin reporter was a good proxy for testing the sensor circuit response function and distinguishing the circuit’s activity from stress-induced pigmentation[54]

### Caution when interpreting reporter response based on transient and stable transgenic assays

We used the transient *S. viridis* protoplast transformation as an efficient platform to run the DBTL cycles for synthetic biology designs before moving to the more laborious process of stable transformation in a homologous system. We evaluated the reporter expression kinetics and dynamics of the pOp6/LHGR-N system with two different promoters for the LHGRN expression using time course imaging of luciferase (Fig. 2A). Strong expression of LHGRN by *pZmUbi1* promoter coincided in the maximum induction of reporter and basal level expression in both protoplasts and stable transgenics (Fig 3-4). At first, the higher fold induction of the reporter and lower background in protoplasts promoted suggested the *pZmEF1a* promoter was an ideal driver for the LHGR-N/pOp6 system (Fig. 2A-B). However, the stable transgenic line pSENT162 harboring *pZmEF1a::LHGR-N* showed weak bioluminescence and anthocyanin by DEX application (Fig S5). This discrepancy between the transient assay and the stable transgenics could result from the high number of plasmids in the protoplast system versus the low-copy to single integration of transgenes in the stable transgenic lines. Alternatively, higher background levels of reporter expression in protoplasts could be caused by insufficient sequestration of LHGRN proteins by cellular chaperones or changes in cellular chaperone activity as the result of stress due to protoplasting [55]. Thus, caution must be taken when extrapolating results observed in protoplasts to the activity of constructs once stably integrated into the genome.

### HSI detection and detection limits

We used ARI and the difference between two wavelengths (specifically, 550 nm and 700 nm) to evaluate and estimate anthocyanin content [51]. Some disadvantages to the ARI approach are that it leverages only two wavelengths (and ignores any potential detection signal in neighboring wavelengths) and does not consider the impact of background spectral variation occurring in the 550 and 700 nm wavelengths. MI-ACE overcomes these disadvantages by leveraging the entire spectral range and learning a discriminative target concept that places a more significant weight on wavelengths most informative for detection in a given background setting. The potential advantage of MI-ACE over ARI is that, given training data representing the expected background and target variations, detection performance can be optimized and enhanced over a simple spectral index. The method allows the use of all discriminative information throughout the hyperspectral signature collected (as opposed to the fixed small number of wavelengths used in ARI), and the optimization tunes the discriminative signature to put more emphasis on the wavelength responses that allow the best differentiation between target and non-target regions. The disadvantages of MI-ACE, however, are that it requires a representative target and background training set that is available to learn the discriminative target concept. The detection performance of MI-ACE will depend on the discriminability between background and target spectral responses and the availability of representative training data. If the environmental conditions and settings deviate from those covered by the training set, MI-ACE performance will likely degrade. Thus, it is essential to consider the environmental conditions the sensor plants may encounter when developing an MI-ACE model before deployment.

### Limitation of chemical diffusion through plant cell wall and epidermal layer

As seen in the limited pattern of bioluminescence and anthocyanin induction in a marginal area of leaves, weak diffusion of steroid hormones through plant tissue can limit systemic induction. The presence of an epicuticular wax layer and rigid cell wall of *Setaria* leaf tissue may be responsible for the low diffusion rate of steroid hormone into leaf tissue, similar to the case of herbicide [56]. Furthermore, the suberin lamellae around the bundle sheath cells of the *Setaria* leaf may pose an additional barrier in the apoplastic transport of solute [57,58]. Thus, the wide use of glucocorticoid receptor-based circuits in some C4 monocot species may be limited due to the limited diffusion of the ligand. More hydrophilic steroids, such as DSP and TA, facilitate its penetration into the leaf tissue, resulting in rapid and substantial reporter expression, but induction was still localized (Figure 5). Since the uptake of specific chemicals varies significantly between plant species, developmental stages, and environmental conditions [59], this reporter system would be a great choice to track the permeability of chemicals with various formulae, such as application method and ratio of chemical and surfactant/adjuvant throughout whole plant and life cycle. To ensure the systemic induction of a target gene, screening small molecules with a high diffusion rate across the leaf epidermal barrier should be coupled with developing chemically inducible plant sensor circuits. One promising inducible system could be based on the PYR1-HAB sensor circuits by selecting new sensors that bind to new ligands with high penetrance into various plant species and tissues [3,60]. Such systems could be used to develop plant-based sensors to monitor chemical exposure and increase plant and human health in agricultural settings.

## Materials and Methods

### Generation of constructs

All constructs used in this study were built with the Golden Gate cloning system [17,18] using Thermo FastDigest restriction enzymes and T4 DNA Ligase (Thermo Fisher Scientific, Waltham,USA). Many L0 regulatory sequences and some L1 modules were repurposed from prior studies [61]. All domesticated and synthesized L0 module insert sequences and position of L1 backbone are summarized in **Supplementary table 1**.

For L1 and L2 constructs, layout and modules of lower order are also presented in **Supplementary table 1.** All L0 modules in this study are deposited in the addgene (https://www.addgene.org/depositing/83475/) and their addgene IDs and identifier #s are noted in the same **Table S1**. Schematics for the multigene construct using 2A skipping peptide is illustrated in **Supplementary** Figure 7. Assembly reactions were performed on a thermocycler with the following steps: 2 minutes incubation at 37℃, 3 minutes at 16 ℃, both steps repeated 20∼30 times, followed by incubation for 5 minutes at 50℃ and 10 minutes at 80℃. To improve the recovery of assemblies with C2 modules, the 2nd annealing step in the assembly reaction was adjusted from 16 ℃ to 10℃.

### Generation of *Setaria* transgenics

L2 constructs listed in **Supplementary table 1** were transformed into Agrobacterium strain, *AGL1*. Setaria transformation on *Me034v-1* accession was generated in the Danforth Plant Science Center tissue culture facility using the procedure previously described in [62].

### Plant growth conditions

*Setaria viridis* accessions *Me034v-1* and its derived transgenics were used for most assays. To reduce dormancy, *Setaria* seeds were treated overnight in a 10% Hickory liquid smoke solution (B&G Foods, Ontario, Canada). Then, liquid-smoke treated seeds were stratified in wet soil and stored at 4 ℃ for at least 7 days. After cold treatment, plants were moved into the greenhouse to germinate. Seeds aged over 2 years were directly sowed into wet soil and moved to the greenhouse or growth chamber without liquid smoke and cold treatment.

Plants were grown in a soil mixture of Metromix 360 (Hummert International, Earth City, MO, USA) and Turface MVP (Profile product LLC, Buffalo Grove, IL,USA). Plants were grown at the Donald Danforth Plant Science Center (St.Louis, MO, USA) under greenhouse growth conditions, 28/ 25 ℃ (14 hr day/ 10 hr night), with a supplementary lighting scheme in the range of 800-1000 μmol m ^-2^s^-1.^.

### Transgenic selection based on Taqman copy number analysis

Taqman copy number analysis was performed to evaluate the number of transgenes at the T0 and transgene segregation in the following generation. Seedling leaves from each line were harvested into a 96-well plate. Genomic DNA extraction and taqman qPCR with 2 technical repeats were performed at Cicadea biotech (St.louis, MO, USA). Taqman copy number analysis was performed as previously described in [61], except for the setaria reference probe sets (5’-AGGCGCATACGTTACCTATA C-3’, 5’-CATGAGGCTGGAACGAAATCT A-3’ and /5HEX/TGCATTAGA/ZEN/TCGAACACCAGCTCGT/3IABkFQ/), designed on *Setaria Leafy* gene, *Seita.7G222300*.

### Setaria mesophyll protoplast transformation

Isolation of mesophyll protoplast from *Setaria* leaves and Polyethylene Glycol (PEG)-mediated transformation were developed after major modifications from [63]. The latest emerged leaves from 10-15 day-old *S. viridis* seedlings were harvested and chopped into leaf strips with a fresh razor. 1∼1.5 g of cut leaves are resuspended in 20 ml of enzyme solution (1.5% Cellulase R10 (Yakult, Tokyo Japan), 0.3% Macerozyme R10 (Yakult, Tokyo, Japan), 10 mM MES (pH 5.7), 0.4 M Mannitol, 1 mM MgCl_2_, 1 mM CaCl_2_, 5 mM β-mercaptoethanol, 0.1% BSA and 50 ug/ml Carbenicillin) for about 4 hours with gentle shaking (10∼20 rpm) at RT in the dark. After cell wall digestion, protoplasts are released by shaking at 90 rpm for 20 minutes and filtered through a 70 μm cell strainer to remove cell debris. After centrifugation at 200 x g for 5 min, the pellet is resuspended into 8 ml W5 buffer (154 mM NaCl, 125 mM CaCl2, 5 mM KCl, 2 mM MES-KOH (pH5.7). Resuspended protoplasts were overlaid onto a 5 ml 0.55 M sucrose solution and centrifuged in a swing bucket rotor at 500 x g for 5 minutes without brake. The protoplast interfacial layer is recovered and washed with W5 buffer. After resuspension with W5 buffer, the protoplast yield was measured using a hemocytometer. Protoplasts were pelleted at 200 x g for 5 minutes and resuspended in an appropriate volume of MaMg buffer (0.4 M Mannitol, 4 mM MES-KOH (pH5.7), 15 mM MgCl_2_) to give a final density of 1 x 10^6^ cells/ml. For each transformation, 2 x10^5^ cells were mixed with 10 μg of each plasmid and then an equal volume of a 40% PEG/CalCl_2_ solution (40% PEG 4000, 0.2 M Mannitol, 100 mM CaCl_2_) was added to the protoplast-plasmids mixtures. Mix the tubes gently by inversion and incubate the transformation mixture in the dark for 20 minutes with gentle shaking at 20∼30 rpm. The reaction was stopped by adding a 2 x volume of W5 buffer and harvesting the protoplasts by centrifugation at 200 x g for 5 minutes. Protoplasts were washed with W5 buffer and resuspended with 1 ml of W5 buffer with 5% fetal bovine serum, 500 µg/ml carbenicillin for downstream assays. For bioluminescence imaging,1 transformant was supplied with 1 mM luciferin and split into 4 wells in a black 96-well culture plate (Greiner bio-one, USA).

### Bioluminescence imaging

To evaluate the bioluminescence induction from transgenic tissue containing luciferase reporter constructs, 9 mm in diameter leaf punches from different lines were arrayed in a black 96 well-plate (Greiner bio-one, USA). Each well was filled with 150 μl of 5 mM luciferin solution with 0.001% BREAK-THRU^®^ OE446 (Evonik, Essen, Germany) surfactant and 500 μg/ml carbenicillin. The imaging plate was transferred to a dark chamber set to 22 ℃. Luciferase recording and analysis were performed as described previously, without any lighting [64,65]. For chemical sensor constructs, ligands are added directly to the luciferin solution at specific time points, as indicated in the figures.

### Chemical application

Dexamethasone (CAS# 50-02-2), Triamcinolone acetonide (CAS# &6-25-5) and Dexamethasone 21-phosphate disodium salt (DSP, CAS# 2392-30-4) were purchased from Millipore Sigma-Aldrich (St.Louis, USA).

For the ultrasonic mist-fogger, commercial ultrasonic diffusers were used within a tight-sealing container placed inside a fume-hood. The appropriate concentration of ligands diluted into water with 0.001% BREAK-THRU® OE 446 surfactant (Evonik Industries AG, Essen, Germany) were poured into the diffuser reservoir to give a final concentration of ligand/volume in the sealed container. Plants in pots and diffusers were placed into the container inside the fume hood until the diffuser evaporated the ligand containing water to saturate the sealed container overnight. The next day, the container was opened to let the surface-coated chemical air-dry for at least 10 min before the treated plants were returned to the growth space until observation.

### Anthocyanin extraction protocol

Leaf anthocyanin extraction was performed as described in [66], with the following modifications. At least 20 mg of leaf tissue was harvested in a 2 ml round-bottomed microfuge tube with two 3.8mm stainless beads, flash frozen in liquid nitrogen (LiqN_2_), and stored in -80 ℃ until processed. Frozen tissue was ground under LiqN_2_ in a MM400 Mixer Mill (Retsch, Dusseldorf, Germany). After lyophilization, 300 μL of methanol with 1% HCl was added to each tube and then incubated overnight in a dark refrigerator (4℃). Then, each tube was brought up to 500 μl by adding 200 μl deionized H_2_O. 500 μL chloroform was added, mixed, then the tubes were spun in a microfuge at 21,000 x g for 5 minutes. 400 μl of clarified supernatant was transferred to a new tube, and then 400 μl of a 60% Methanol/59% H_2_O/1% HCl solution was added to each tube. Anthocyanin content (Anthocyanin/mg fresh weight) was obtained by reading at 530 nm and 657 nm on a SpectraMax® M3 plate reader (Molecular Devices, San Jose, USA) or a spectrophotometer, and the following calculation was used (A530 - A657)*1000/ mg fresh weight.

### Western blotting

10 mg of leaf blades was frozen with liquid nitrogen and grounded with two stainless beads on a MM400 Mixer Mill (Retsch, Dusseldorf, Germany). Lyophilized tissues were resuspended into the 200ul of 2x laemmli sample buffer with the 5% β-mercaptoethanol and boiled at 95℃ for 5 minutes. After centrifugation, 10 ∼ 20 ul of each supernatant was used for SDS-PAGE followed by Western blotting using anti-FLAG M2 antibody (Sigma-Aldrich, USA)

### Plant materials for hyperspectral imaging

For imaging scans containing plants constitutively expressing SvC1 and SvR1 (pSENT98) and ME034v-1, a total of five and three biological replicates were used respectively. In the case of the inducible lines pSENT166 (#6-7 and #8-7) and null plants (pSENT166 #8-10) sprayed with DSP, TA and Dex, a total of four biological replicates were used for imaging. All plants were imaged 14-15 days after planting.

### Hyperspectral Development Platform (HDP) and Scans

The HDP includes a high-resolution VNIR camera (Headwall Series E, 400 to 1000 nm, with 923 spectral bands) integrated on a Kuka KR Quantec Pro robotic arm. The HDP has dedicated halogen illumination from two angles, and top-view images of taped leaves were gathered. We used a mate gray vivid PVC board (Palight^®^) as the image background to reduce the reflectance from the background itself.

Before images were taken, white reference and dark reference images were captured to calibrate the raw hyperspectral image data cube into reflectance values. For hyperspectral image scans, *Setaria viridis* (*S. viridis*) plants were lined up so the plants were not touching. Top view hyperspectral images were acquired from the plants by using the HDP at 30 cm from the samples.

### PlantCV for Hyperspectral Image Segmentation and ARI Index Calculation

PlantCV is an open-source package of tools aimed at flexible plant phenotyping, including the analysis of hyperspectral images [48,49]. A number of open-source tools have been added to PlantCV in order to analyze hyperspectral data in a reproducible and consistent manner. Specifically, modules **analyze_spectral.py** and **analyze_index.py** (https://github.com/danforthcenter/plantcv/blob/main/plantcv/plantcv/analyze/spectral_in dex.py) were used. Tutorials for hyperspectral image analysis have been developed to extract and quantify important information https://plantcv.readthedocs.io/en/stable/tutorials/hyperspectral_tutorial/. The hyperspectral sub-package of PlantCV contains many indices which are available depending on the wavelengths captured by the particular system. For early anthocyanin detection with PlantCV, soil-adjusted spectral index (SAVI) [67] was used to segment plant tissue from the background, resulting in a binary mask that identifies plant material. Once the plant material was identified, manually a custom polygon region-of-interest (ROI) was created for each plant within an image https://plantcv.readthedocs.io/en/stable/tutorials/roi_tutorial/#tutorial-region-of-interest-tutorial. The Anthocyanin Reflectance Index (ARI) was calculated using the reflectance of the segmented plant pixels.

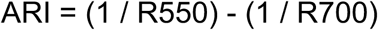

The ARI [33] of each plant was calculated for **pcv.hyperspectral.analyze_index** function. The proportion of pixels was obtained and was plotted with the index reflectance value. A Kolmogorov-Smirnov (KS test) was calculated to compare the sample against the respective control in order to quantify the distance between their distributions [68].

### MI-ACE analysis

The MI-ACE detection algorithm was applied in order to distinguish between the treated and untreated plant specimens on a sub-pixel level. MI-ACE is designed to estimate a discriminative target signature from training data with imprecise (multiple instance) labels. MI-ACE analysis begins first with a preprocessing stage, and followed by an analysis stage. Test and training sets are then iterated, with each image section being left-out as a test set once. In training data, image sections are all labeled with either target absent (background only, negative) or target present (background + scaled target signature **s**, positive) under Gaussian noise assumptions. Then, background mean 𝜇_*b*_ and covariance 𝛴_*b*_ are computed from the negative bags. The data is whitened and normalized. At this point, the method initializes the discriminative target concept, ***s***, randomly to serve as an initial point. The analysis notebook is then ready to perform MI-ACE on the whitened and normalized data. It begins iterating through the data, maximizing the ACE statistic on positive bags ***j*** and minimizing it on negative bags ***i***. More specifically, it selects instance ***x’’*j*** from each positive bag ***j*** with maximum ACE(***x’’, s***), and instance ***x’’*i*** from each negative bag ***i*** with minimum ACE(***x’’, s***). The target signature is then updated as a series sum of these products per the equations:

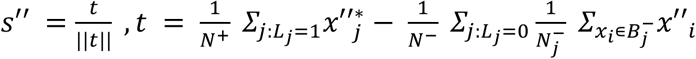

This resulting signature is normalized and compared with the previous iteration of ***s***. If the two are equal, then the algorithm has converged and the final signature is outputted. Otherwise, it will continue iterating until convergence is reached.

### Preprocessing and Mask Generation

In order to run MI-ACE on the plant dataset, positive and negative masks were generated for the treated and untreated plants using the plantCV library. The **pcv.roi_objects** function is used to detect and extract objects representing each plant given a specified region of interest (ROI). These masks are then used to generate positive and negative bags, which are in turn applied to the MI-ACE detection. To begin pre-processing, the original hyperspectral image was segmented into a grid of 2 number image sections (positive for anthocyanin and negative for control).

### MIACE Algorithmic Experimental Design

Image data was split into “bags”, or subsections, based on anthocyanin presence in order to determine a target signature for subjects with anthocyanin present. The training data was split into its corresponding bags by visual inspection and division of image sections based on known locations of the anthocyanin presence. Bags were labeled as positive (contains at least one target data point) or negative (no target points). MI-ACE then optimized the Adaptive Cosine Estimator (ACE) detection statistic on this training data. The ACE detection statistic is iterated multiple times until it converges to an estimated target signature [34].

## Supporting information

Supplemental Table 1 Addgene Plasmids

## Acknowledgements

We would like to thank Dr. R. Keith Slotkin and Dr. Kaushik Panda for helpful discussion on transposon insertion in *SiC1 gene* and Dr. Veena Veena and Todd Finley for transformation of *S.viridis*, Me034v-1. This work was supported by Defense Advanced Research Projects Agency Advanced Plant Technologies (DARPA-APT, HR001118C01327). The views, opinions, and /or findings expressed are those of the authors and should not be interpreted as representing the official views or policies of the Department of Defense of the U.S. Governments.

## Supplementary Figure and Legends

**Supplemental Figure 1.**
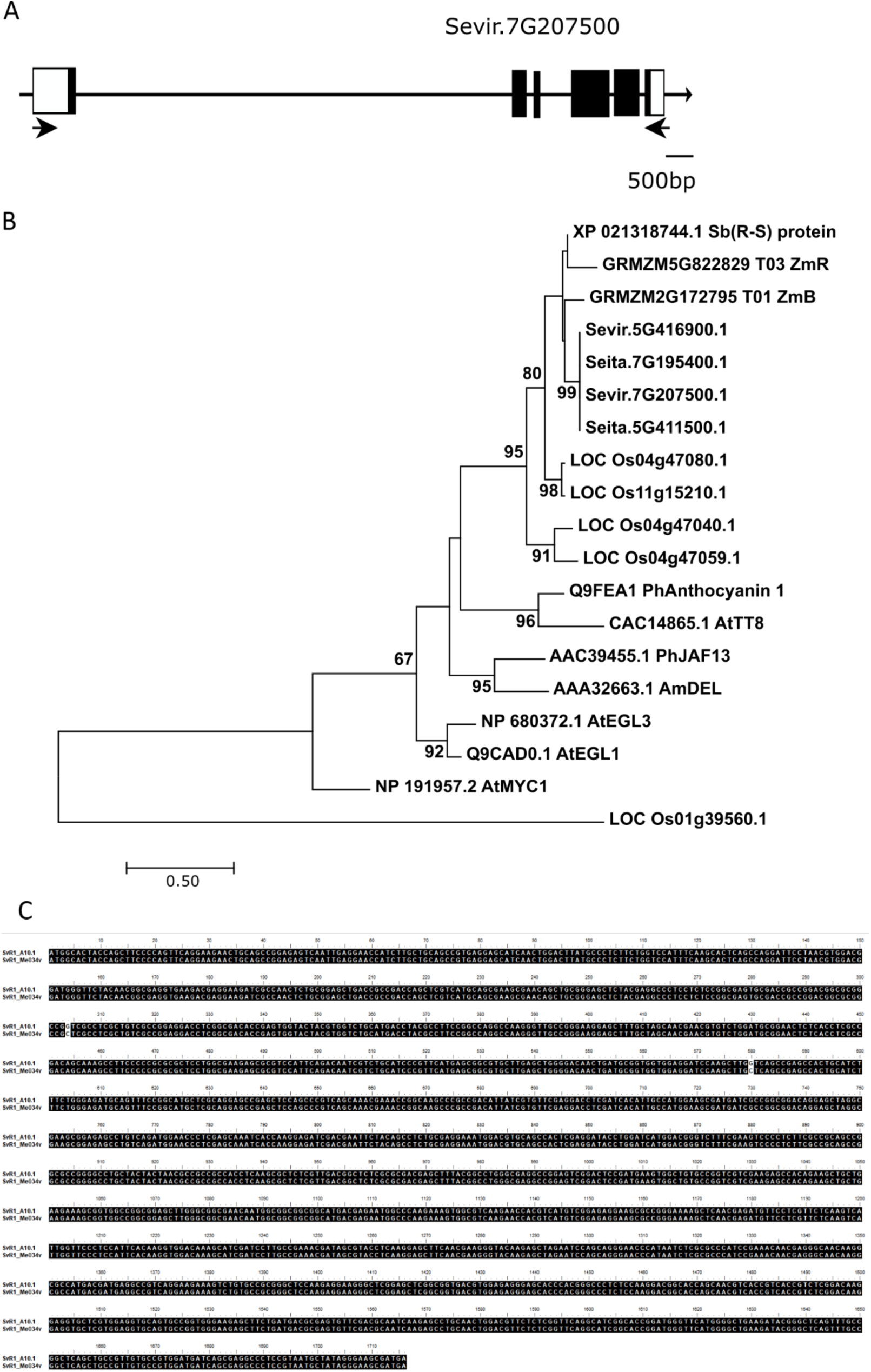
Phylogenetic analysis and cloning of R1 homologs. (A) Structure of *SvR1* (Sevir.7G207500) gene in *Setaria viridis A10.1* genome. Filled boxes and empty boxes indicate the exon and UTR regions, respectively. Arrows represent the location of primers for the full-length cDNA isolation. Scale bar= 500 bp. (B) Molecular phylogenetic tree of the myogenic bHLH transcription factors in the *Setaria*, rice and maize associated with ZmR by Maximum Likelihood. The length of the branches represents the evolutionary distance between ancestor to descendent nods. The numbers represent the confidence level of the specific branch. Abbreviation in tree as follows: ZmB, *Zea mays Boost*; Sb (R-S), *Sorghum bicolor R-S*; PhAnthocyanin1, *Petunia Hybrid Anthocyanin1* (AN1); AtTT8, *Arabidopsis thaliana Transparent testa* (*TT8*); PhJAF13, *Petunia hybrid JAF13*; AmDEL, *Antirrhinum majus Delia*; AtEGL3, *Arabidopsis thaliana Enhancer of Glabra 3.* (C) Nucleotide alignment of SvR1 open reading frame (ORF) between A10.1 and Me034v-1. putative SvR1 cDNA sequences were deduced from A10.1 reference genome and isolated cDNA from the leaf tissue of Me034v-1.

**Supplemental Figure 2.**
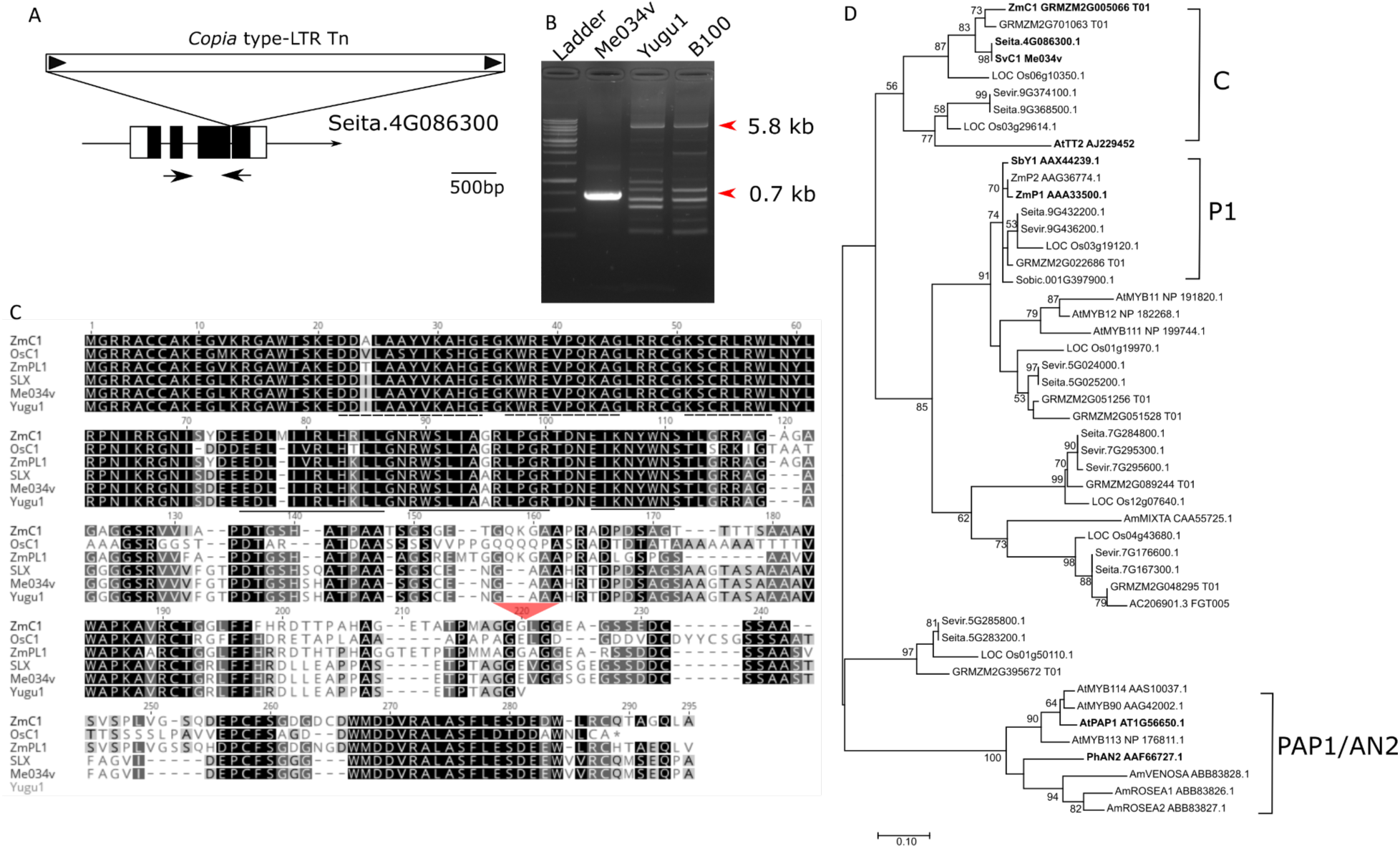
Phylogeny analysis of C1 homologs and identification of a transposon insertion in *S. italica* C1 gene. (A) Gene structure of C1 gene in *Setaria viridis* and transposon insertion in *italica* cultivars, *Yugu1*. Arrows represent the location of primers used for the presence of Copia type transposon in Fig S2B. (B) Presence of Copia-type LTR transposon in C1 gene of *S.italica*, *Yugu1* and *B100* accessions. PCR was performed with the primers (Fig S2A, Online Methods). From the left, 1 kb ladder (Gold bio), Me034v-1, Yugu1 and B100. Expected bands with/without copia-transposon are noted as red arrowheads. (C) Amino acid sequence alignment of C1 coding sequence from Yugu1, SLX and Me034v-1 with the homologs in monocot. Conserved α-helics structures in two MYB repeats are shown as underlined (R3) and dashed lined (R2). Red triangle indicates the insertion point of copia type LTR transposon on C1 gene in two *italica* accessions. *ZmC1*, *Zea mays C1; OsC1, Oryza sativa C1; ZmPL1, Zea mays Purple plant1.* (D) Molecular phylogenetic tree of the R2R3Myb family in the Setaria and rice and maize by Maximum Likelihood. The length of the branches represents the evolutionary distance between ancestor to descendent nods. The numbers represent the confidence level of the specific branch. Separated clades were noted as brackets on the right side as C, P1 and PAP1/AN2.

**Supplemental Figure 3.**
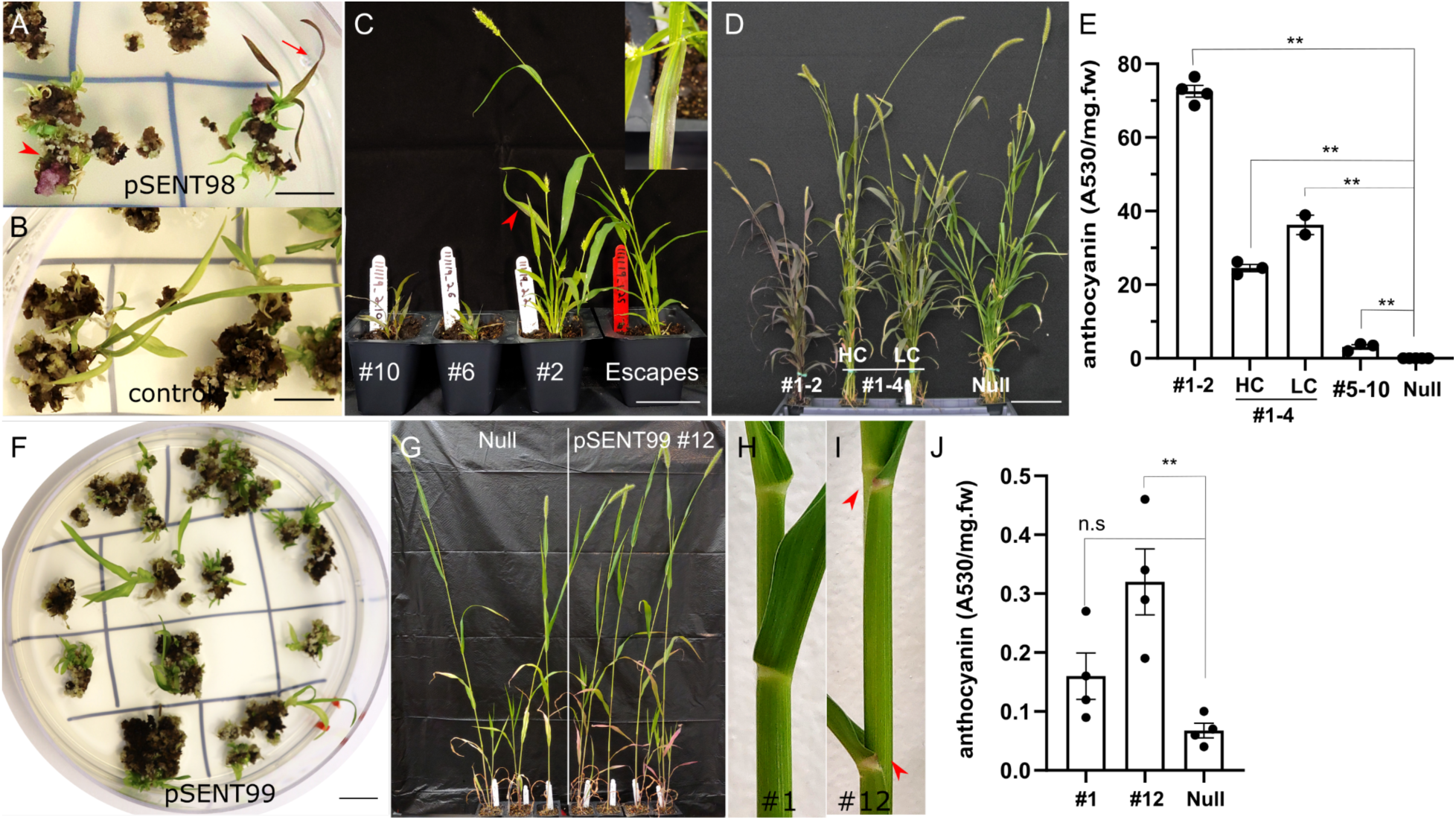
Characterization of transgenic coexpressing SvC1 and SvR1 in *S.viridis*. (A) Purple callus (red arrow heads) and purple shoots (red arrows) of callus transformed with pSENT98 (pZmUbi1::SvR1-E2A-SvC1) on regeneration media. (B) Typical green shoots and greening calli with the control GUS construct during regeneration. Scale bar = 1 cm. (C) Pleiotropic phenotypes from primary transgenic pSENT98. # denotes the number of independent events from tissue culture. Escapes means non-transgenic among regenerates. Single insertional transgenic events, #10 and #6 showed the strong anthocyanin accumulation but stunted in growth, leading to the absolute sterility. #2 showed the chimeric anthocyanin sectors (inset figure) and its anthocyanin phenotype was not inherited in the subsequent generations. Scale bar = 5 cm. (D) purple plant in T1 generation of pSENT98 #1. Multi-copy transgene event #1 was segregated into the homozygous fixed individuals (#1-2), various copy number of transgenic line (#1-4 as representative) and null siblings. T2 lines from #1-4 were segregating into the high copy transgenic (# of transgene >5), low copy transgenic (number of transgene=4) and null. Scale bar = 10 cm. (E) Anthocyanin extraction from T2 lines of pSENT98. 2 fully emerged leaves from 10 days old seedling of fixed #1-2, segregating lines from #1-4, #5-10 (single insertional transgenic homozygotes but low anthocyanin pigmentation) and null siblings were harvested for extraction. Each bar and vertical line indicated the mean and standard error. Statistical significance denotes as ** (P<0.01, two-tailed T-test). (F) Green shoots and calli from pSENT99 (pOsUbi3::SvR1-E2A-SvC1) regeneration. Scale bar = 1 cm (G) anthocyanin expression of senescensing leaves in pSENT99 #12. Segregating nulls siblings (left), homozygotes (right). Leaves of transgenic were turning pink as senescence progressed but not in the null. (H-I) Anthocyanin accumulation (red arrowhead) in the ligule of pSENT99 #12(I) but not in pSENT99 #1 (H) line showed non-pigmentation like wild-type Me034v-1. (J) anthocyanin contents of leaves from pSENT99 #1, #12 and null seedlings. 4 individuals from #1 and #12 T2 homozygous lines were harvested for extraction. Statistical significance denotes as ** (P<0.01, two-tailed T-test) and n.s (not significant).

**Supplemental Figure 4.**
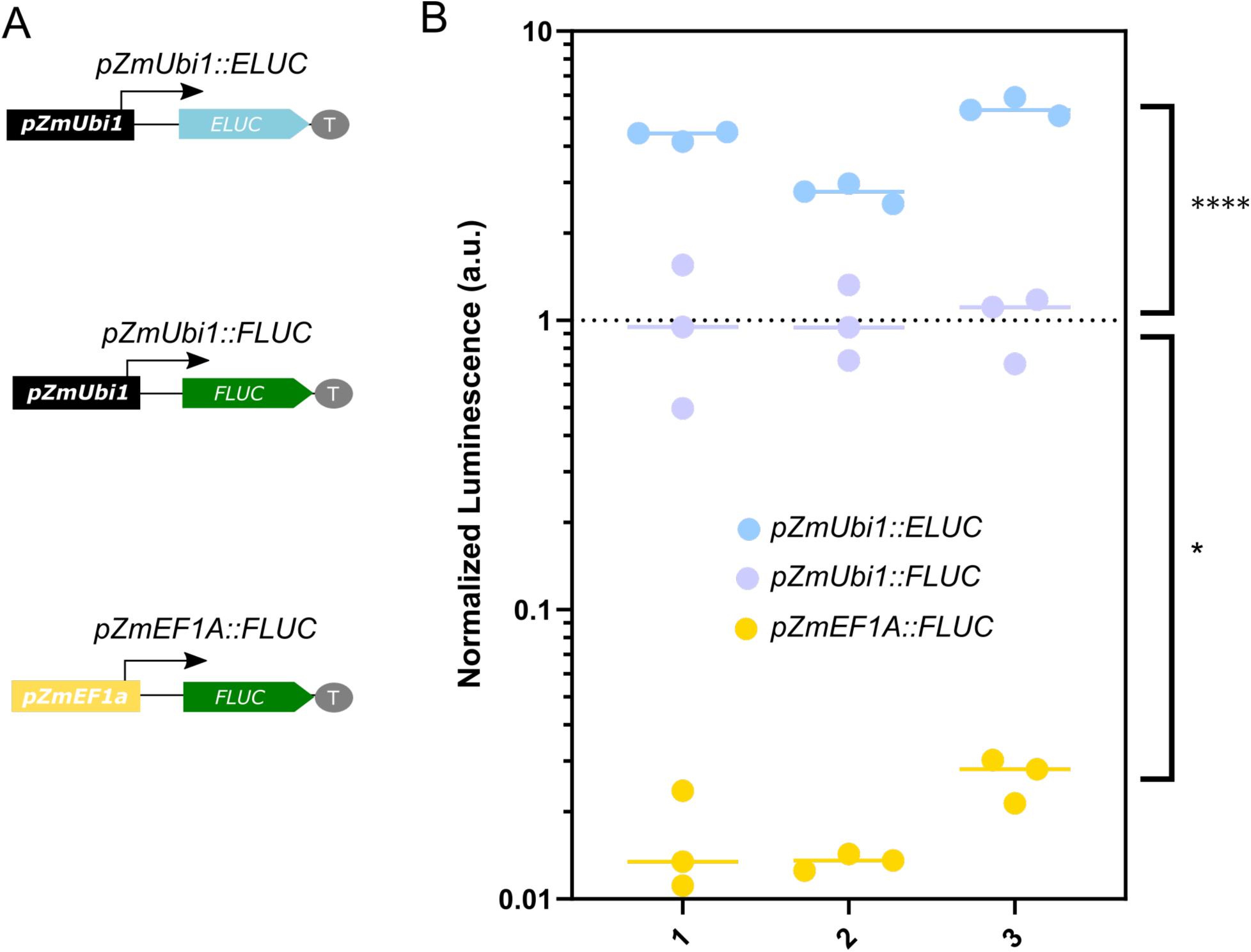
Comparison of *ZmUbi1* promoter and *ZmEF1a* promoter strength and the catalytic activity difference between ELUC and FLUC in *S. viridis* protoplasts. (A) Construct scheme of promoter luciferase construct. *pZmUbi1*; *Zea mays Ubiquitin 1* promoter, ELUC; Human codon optimized *Pyrearinus termitilluminans* larval click beetle luciferase, FLUC; *Photinus pyralis* (Firefly) Luciferase, *pZmEF1a; Zea mays Elongation factor 1a* promoter. Arrow represents the direction of promoter in downstream gene transcription. (B) Normalized expression level of promoter luciferase construct in the mesophyll protoplast of *S.viridis*. lines and dots represent median and individual bioluminescence, respectively, from transfection with representing constructs after normalization to the mean luminescence of *pZmUbi1::FLUC* within the same batch of protoplast (X axis). P-values were calculated using the nested one-way ANOVA analysis, *P≤ 0.05, **** P≤0.0001.

**Supplemental Figure 5.**
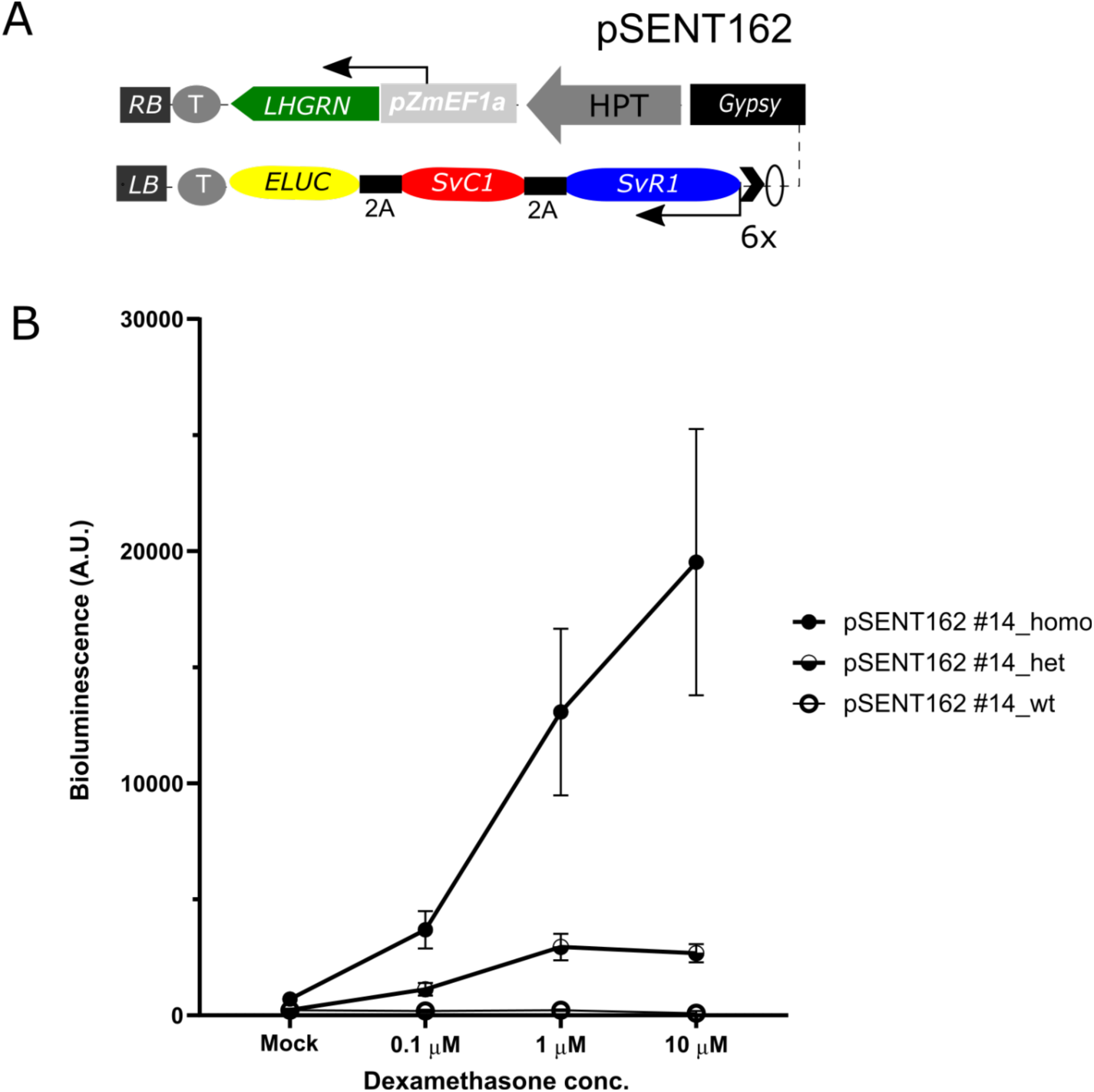
Ligand and gene dosage dependent induction of pSENT162 in *S. viridis.* (A) construct layout of pSENT162 dexamethasone inducible with pZmEF1a::LHGR-N. Gypsy, Drosophila Gypsy insulator; 2A, 2A-skipping peptide; ELuc, Human codon optimized *Pyrearinus termitilluminans* larval click beetle luciferase; RB, Right border of transfer DNA (T-DNA); LB, Left border of t-DNA; black chevron, minimal 35S promoter; O, Lac operator sequence. (B) bioluminescence induction of leaf discs from pSENT162 #14 T2 lines by 3 different doses of dexamethasone and mock (0.1 % Methanol). Filled, semi-filled and empty circle represents homozygotes, heterozygotes and nulls, respectively. Vertical line of each point indicates the standard error from at least 4 bio-reps (individual T2 plants).

**Supplementary Figure 6.**
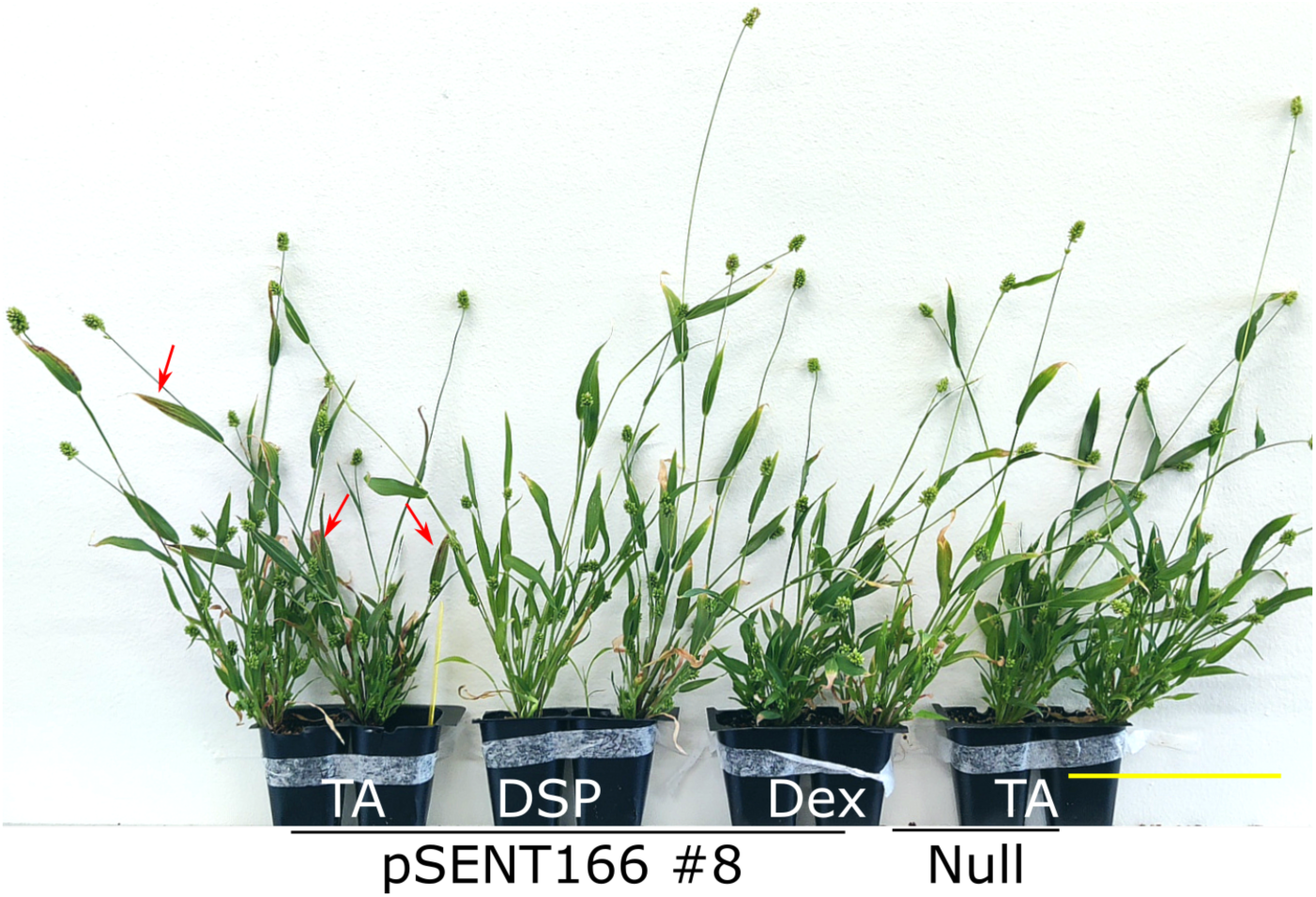
Systemic induction of the anthocyanin reporter of pSENT166 by DEX analog, Triamcinolone Acetonide (TA). comparison of systemic induction of reporters by various steroids. 10 days old pSENT166 #8 and their null were treated with 1 μM TA, DSP and Dex by ultrasonic fogging. Images were taken at 32 days after steroid application. Red arrows indicate the red-pigmented region in flag leaves. Scale bar =5 cm

**Supplementary Figure 7.**
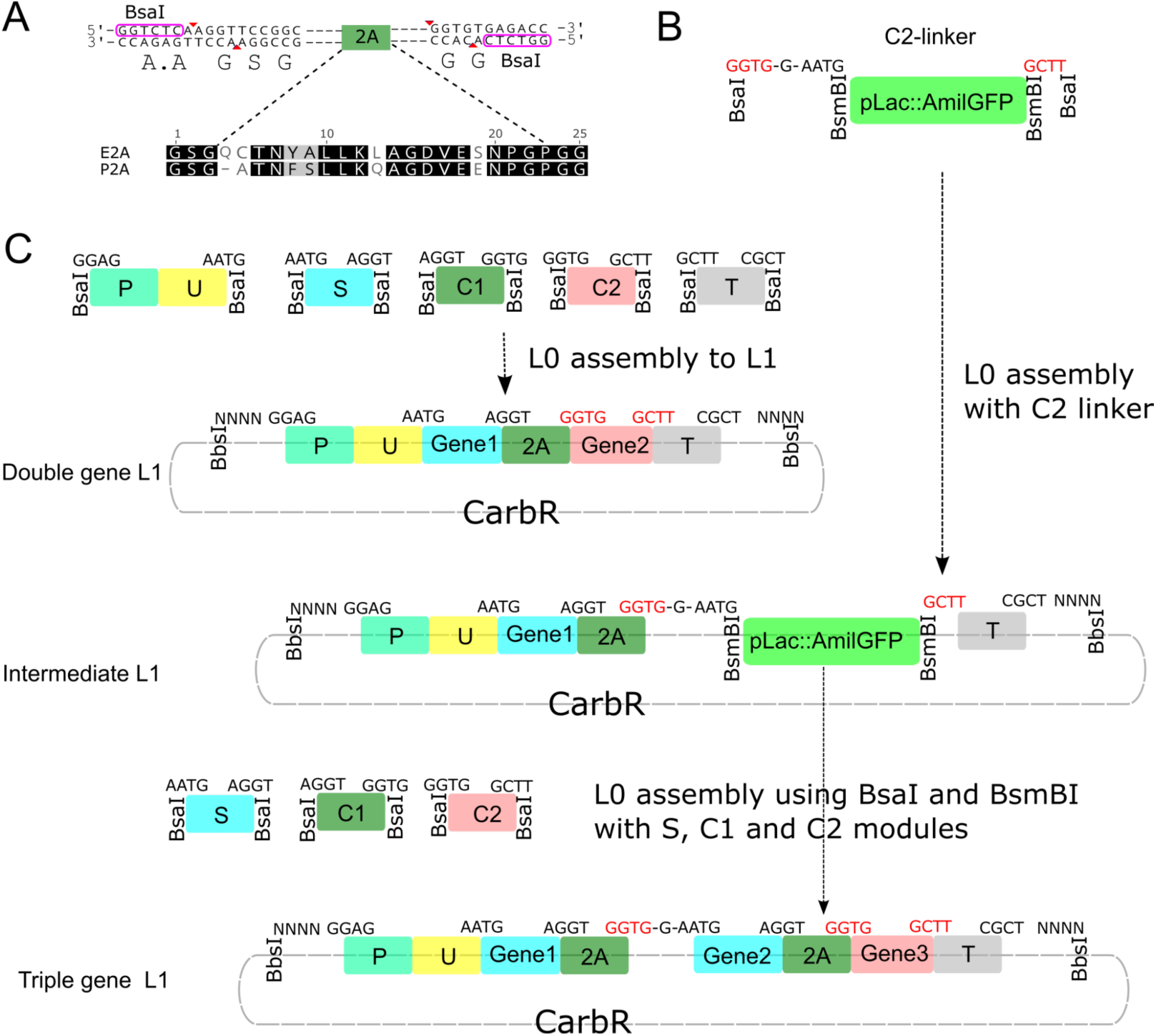
Schematic representation of multi-gene construction using 2A peptides in the golden gate system. (A) Designs of 2A peptide sequence in L0-C1 position. Glycine-serine-glycine (GSG) linker and GG-linker are added N’-terminus and C’-terminus of 2A peptide, respectively, for the efficient peptide cleavage and cloning purpose. E2A, equine rhinitis a virus 2A; P2A, porcine teschovirus-1 2A. Nucleotide sequence flanking 2A peptide is shown at top and amino acid sequence of 2A peptide is shown at the bottom. BsaI recognition sequences are marked with purple ellipses and red triangles show the position of restriction sites. (B) Design of C2-linker module for multi-gene cloning. Color selection marker *pLac::AmilGFP* was placed between two Type II S restriction sites BsaI and BsmBI and additional sequences for translational fusion of further cloning of SC module *in-frame* [69] (L0-23, Supplementary table 1). Bright yellow color/green fluorescence with the presence of pLac::AmilGFP can be used for the screening of intermediate L1 construct cloning and subsequent cloning. (C) Examples of L0 assemblies to build the multi-genic constructs. Top 5 box represents 5 standard L0 modules flanked with BsaI restriction sites and 4 bp fusion sites above boxes. P, promoter; U, 5’UTR; S, signal sequence; C1, Coding sequence 1; C2, Coding sequence 2; T, Terminator. The C1 position of 2A peptide was positioned between two independent genes in the S and C2 position to make a double-gene construct. For the multigenic reporters, C2-linker was placed at the L0-C2 position to make intermediated L1 in the first L0 assembly. C2-linker opens to clone additional genes with intervening 2A peptide using BsmBI and BsaI restriction enzymes in the 2nd assembly. CarbR, beta-lactamase resistance; NNNN, 4bp fusion sites which vary on the position of L1 constructs.

**Supplemental Figure 8.**
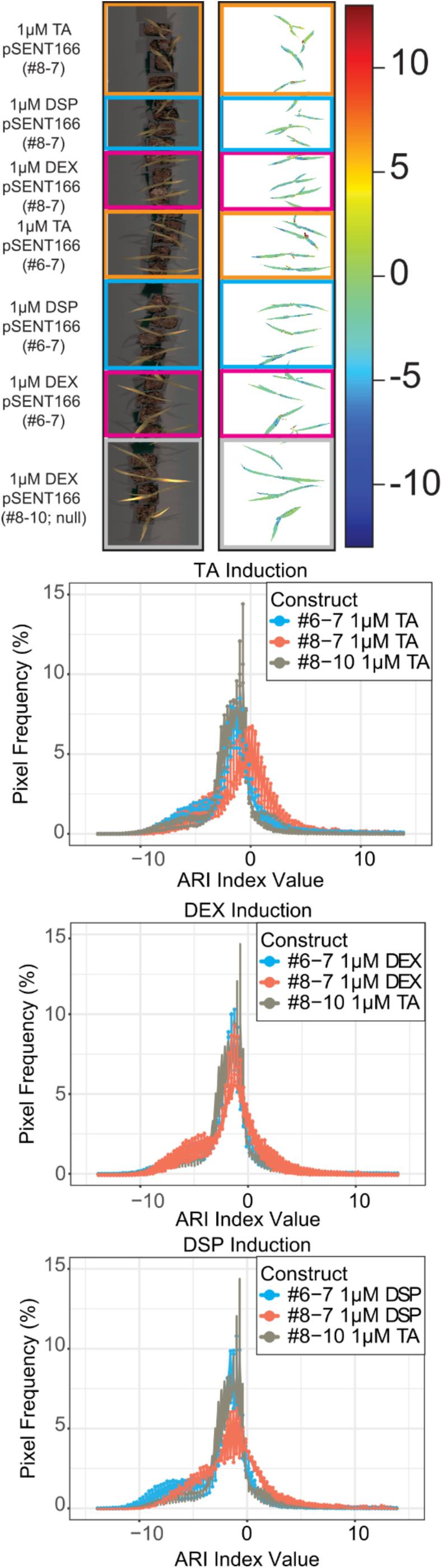
Detection of anthocyanin production in inducible lines pSENT166 (#6-7 and #8-7) and null line pSENT166 (#8-10). A) Pseudo RGB and pseudo-colored Anthocyanin reflectance indices (ARI) from pSENT166 (#6-7 and #8-7, and null line #8-10) *setaria* plants sprayed with 1µM DSP, TA, or DEX. B) Line graphs represent the average ARI index value, n=4. No statistical difference was found in the shift of reflectances.

**Supplemental Figure 9.**
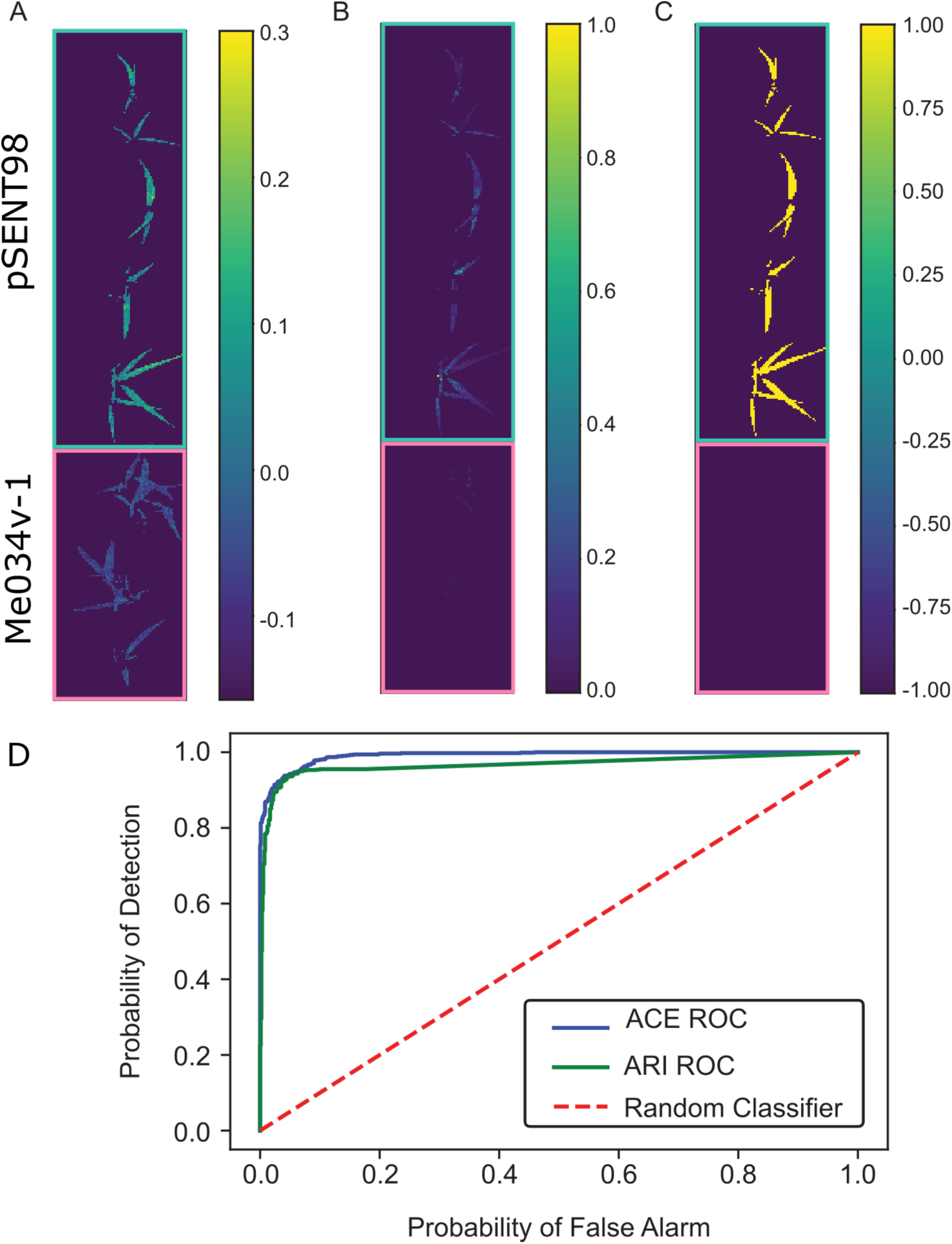
(A-C) Application of Multiple Instance Adaptive Cosine Estimator (MIACE) to detect anthocyanin content in pSENT98 and WT, ME034v-1. (A) MIACE confidence map, (B) Anthocyanin reflectance indices (ARI) confidence map, (C) Truth map (shows where anthocyanin was present, bright yellow means where anthocyanin is present, dark purple represents where anthocyanin is not present and blue corresponds to background. D) Receiver operating characteristic (ROC) curve for both ARI and MIACE approaches, MIACE shows a lower probability of false alarm and higher probability of detection.

## Additional information

**Supplementary Table 1**.Golden gate construct assembly and sequence information

## Online Methods

### Molecular Phylogenetic analysis

To search the homologs of ZmR1 and ZmC1, TBLASTN searches of protein sequence have been conducted on the *Setaria italica* and *Setaria viridis* genome pseudo molecules (https://phytozome-next.jgi.doe.gov/). For the molecular phylogenetic analysis, amino acid sequences of R1 and C1 related genes were downloaded from NCBI (https://www.ncbi.nlm.nih.gov/) and TAIR (https://www.arabidopsis.org/) and their accession number are presented in the Figures S1B and S2D, respectively. Full-length protein sequences within the same gene clusters are aligned using Clustal W with Bootstrap NJ Tree (n=1000)(1) in Bioedit (2). Maximum Likelihood phylogenetic trees were generated by the JTT matrix-based model (3) in MEGA 7(4) with Bootstrap support (n=1000).

### Isolation of SvR1 full-length cDNA and PCR analysis

To isolate the full-length *SvR1* (*Sevir.7G207500*) cDNA, RNA from prior study (5) was used for cDNA synthesis using Improm-II^TM^ reverse transcriptase (Promega, Madison, USA) and Oligo(dT)15 primer (promega) according to the manufacturer’s instructions. cDNA was used for the PCR analysis using Phusion^TM^ HIgh-Fidelity DNA polymerase (Thermo Fisher Scientific, Waltham,USA) with primers (5’-CCTCGTAGCTTCGTCTTTGG-3’ and 5’-AACATTGAGCTGCTCCTCATC-3’). The PCR product was subcloned into the pJET1.2/blunt vector (Thermo Fisher Scientific, Waltham,USA).

To check the presence of copia-type transposons, genomic DNA was extracted from Me034v-1, Yugu1 and B100 accessions using the standard CTAB extraction method (6). PCR was performed with primers (5’-AGGACCTCATCATCCGGCTC-3’ and 5-’CCATTCCTCGTCGGACTCGA-3’) using Phusion^TM^ High-Fidelity DNA polymerase (Thermo Fisher), supplemented with 1M betaine to overcome the uneven GC contents biasedness between the genomic C1 region and transposon.

